# Phase separation of +TIP-networks regulates microtubule dynamics

**DOI:** 10.1101/2021.09.13.459419

**Authors:** Julie Miesch, Robert T. Wimbish, Marie-Claire Velluz, Charlotte Aumeier

## Abstract

Regulation of microtubule dynamics is essential for diverse cellular functions, and proteins that bind to dynamic microtubule ends can regulate network dynamics. Here we show that two conserved microtubule end-binding proteins, CLIP-170 and EB3, undergo phase separation and form dense liquid-networks. When CLIP-170 and EB3 act together the multivalency of the network increases, which synergistically increases the amount of protein in the dense phase. *In vitro* and in cells these liquid networks can condense tubulin. *In vitro* in the presence of microtubules, EB3/CLIP-170 phase separation can co-condense tubulin all along the microtubule. At this condition microtubule growth speed increases up to two-fold and depolymerization events are strongly reduced, compared to conditions with phase separation deficient networks. Our data show that phase separated EB3/CLIP-170 networks impact microtubule growth dynamics beyond direct protein-microtubule interactions.

## Introduction

The microtubule cytoskeleton engages in a plethora of cellular processes, from organelle transport to cell division. To do so, the network dynamically modifies its structure in response to external cues and adapts its architecture to specific cellular functions. Microtubules themselves are highly dynamic polymers that can rapidly cycle between phases of polymerization and depolymerization, a characteristic which is critical for cytoskeletal re-organization (reviewed in Brouhard and Rice, 2018). In cells, microtubules polymerize at their plus-end by addition of GTP-tubulin. After GTP-tubulin addition, GTP is gradually hydrolyzed, resulting in a GDP-tubulin shaft behind the tip. Once the stabilizing GTP-tubulin “cap” disappears from the plus-end, the microtubule switches from growing to shrinking, an event termed catastrophe (Howard and Hyman, 2009; Brouhard et al., 2015; Gudimchuk and McIntosh, 2021). Conversely, microtubules can stop shrinking and switch to regrowth, an event termed rescue. The balance between growth and shrinkage is intimately linked to the addition of free tubulin to the growing microtubule (Walker et al., 1988; Voter et al., 1991).

In addition to these intrinsic modes of regulation, microtubule dynamics can be fine-tuned by Plus Tip-Interacting Proteins (+TIPs) (reviewed in Akhmanova and Steinmetz, 2010). +TIPs are functionally independent and structurally diverse microtubule regulators that concentrate at growing microtubule ends while exhibiting a weak affinity for the microtubule shaft. It is generally accepted that the unique localization of +TIPs at microtubule ends results from their specific binding to the GTP-tubulin cap (Akhmanova and Steinmetz, 2015).

Key integrators of +TIP-networks are the End-Binding proteins (EBs), as they autonomously bind to GTP-tubulin at growing microtubule ends and recruit a battery of non-autonomously binding +TIPs (reviewed in Galjart, 2010). Within the EB family, higher eukaryotes express three proteins termed EB1, EB2, and EB3. These proteins increase microtubule plus-end dynamics by promoting catastrophes and increasing growth speed *in vitro* (Bieling et al., 2008; Vitre et al., 2008; Komorova et al., 2009; Montenegro Gouveia et al., 2010).

A key accessory protein recruited to plus ends by the EBs is the Cytoplasmic Linker Protein of 170 kDa (CLIP-170), which increases microtubule rescue frequency and growth speeds (Perez et al., 1999; Arnal et al., 2004; Bieling et al., 2007; Bieling et al., 2008; Komorova et al., 2002, 2005). CLIP-170 consists of a microtubule binding “head” domain in its N-terminus, followed by a central coiled-coil region and a zinc-knuckle domain, hereafter referred to as the “C-terminal region” (Pierre et al., 1992; Pierre et al., 1994; Diamantopolous et al., 1999). Most studies of CLIP-170 function *in vitro* have focused on truncated versions, containing only the monomeric head domain (H1) or the head domain with a small extension that allows dimerization (H2) (Figure 1A); thus, it is unclear how full-length CLIP-170 contributes to microtubule dynamics.

**Figure 1:**
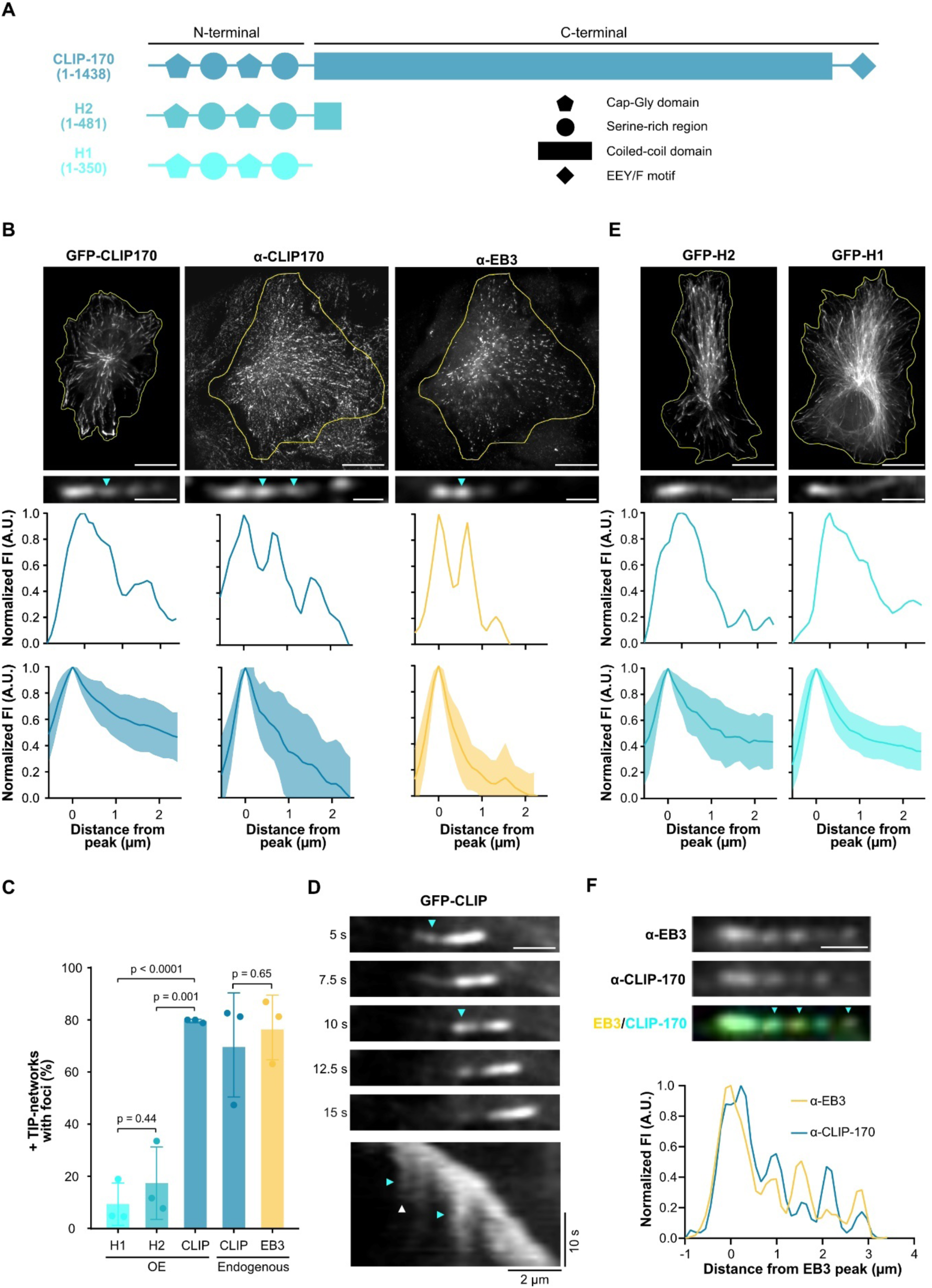
In cells the EB3/CLIP-170 +TIP-network displays liquid properties at microtubule tips. (**A**) Secondary structure of CLIP-170 (1-1438), H2 (1-481) and H1 (1-350) based on (Pierre et al., 1994; Diamantopolous et al., 1999; Goodson et al., 2003). (**B**) Representative images (top) of fixed RPE-1 cells transfected with full length GFP-CLIP-170 or WT RPE-1 cells stained with antibodies to endogenous CLIP-170 and EB3. Representative profiles of +TIP-networks with matching fluorescence line scans (bottom). Cyan arrowheads indicate tailing foci. Scale bars: 20 µm whole-cell, 2 µm insets. Below are quantified mean line scan profiles (dark line) with standard deviation (shaded area) from 3 independent experiments with a total of: GFP-CLIP-170 57 +TIP-networks from 22 cells; anti-CLIP-170 and anti-EB3 58 +TIP-networks from 12 cells for each condition. (**C**) Percentage of +TIP-networks with foci in fixed cells expressing the indicated CLIP constructs or stained with antibodies to endogenous CLIP-170 or EB3 analysis from B and E. Mean with SD from 3 independent experiments. Statistics: one-way ANOVA test. (**D**) Representative time-lapse images (top) and kymograph (below) of +TIP-network from GFP-CLIP-170 expressing RPE-1 cell. Cyan and white arrowheads denote foci formation and dissolving respectively in both time-lapse images and kymograph. Scale bar: 2 µm. (**E**) Representative images of fixed RPE-1 cells transfected with GFP-H2 and GFP-H1. Representative profiles of +TIP-networks with matching fluorescence line scans (bottom). Scale bars: 20 µm whole-cell, 2 µm insets. Below are quantified mean line scan profiles (dark line) with standard deviation (shaded area) from 3 independent experiments with a total of: H2 33 +TIP-networks from 18 cells; H1 47 +TIP-networks from 33 cells. (**F**) Representative +TIP-network from cells stained for endogenous EB3 and CLIP-170 showing partial co-localization of EB3 and CLIP-170 foci (cyan arrowheads) with corresponding fluorescence line scan below. Scale bar: 1 µm.

Early studies of CLIP-170 expression in cells revealed that in addition to its microtubule plus-end localization, it also formed cytoplasmic “patches” that co-localized with EB1 and the dynein-activating protein dynactin (Pierre et al., 1994; Goodson et al., 2003). Based on the physical properties of these CLIP-170 patches, it has recently been suggested that they form by liquid-liquid phase separation (LLPS) (Wu et al., 2021; Jijumon et al., 2022).

LLPS is the process by which molecules spontaneously condense into droplets and de-mix from their surrounding solution, resulting in the co-existence of two unique liquid phases (reviewed in Boeynaems et al., 2018; Hyman et al., 2014; Shin and Brangwynne, 2017). Recently, LLPS has been implicated in driving microtubule-related processes, including spindle assembly (Jiang et al., 2015), nucleation of acentrosomal and branched microtubules (Hernández-Vega et al., 2017; King and Petry, 2020), and centrosome maturation (Woodruff et al., 2017; Jiang et al., 2021). A key shared feature of these processes is the co-condensation of tubulin with LLPS-potent microtubule-associated proteins to catalyze biochemical reactions. Along these lines, condensation of a microtubule +TIP-network has been proposed based on transient multivalent interactions between the +TIPs (Akhmanova and Steinmetz, 2015; Wu et al., 2021). Despite this, LLPS of +TIPs and its function is still unclear.

+TIPs concentrate at the growing GTP-microtubule tip. A combination of *in vivo, in vitro* and *in silico* work has indicated that the GTP-tubulin cap size is 1-60 tubulin layers, indicating a length of < 500 nm (Voter et al., 1991; Walker et al., 1988; Drechsel and Kirschner, 1994; Schek et al., 2007, Seeptapun et al. 2012; Rickman et al., 2017). Puzzlingly, in cells +TIP networks bind to a region at the microtubule end up to 4-fold longer than this, with network lengths up to 2 µm (Komarova et al. 2009; Seepatun et al., 2012; Roth et al. 2019). The discrepancies between +TIP network profiles *in vitro* and in cells could be due to different sizes of the GTP-cap, but also prompts the question of whether network densities and organizational properties could impact network extension.

Interestingly, overexpressed EB3 and CLIP-170 exhibit atypical network formation at the growing microtubule tip. Both proteins leave trailing protein “foci” behind the leading network edge (Mustyatsa et al., 2019; Henrie et al., 2020, Komarova et al. 2009, Nakamura et al. 2012, Mohan et al., 2013). One explanation for these foci is that they form by binding to tubulin remaining in the GTP-state behind the GTP-cap (Perez et al., 1999; de Forges et al., 2016; Henrie et al., 2020). As overexpressed CLIP-170 has the potential to undergo LLPS (Wu et al., 2021; Jijumon et al., 2022), another possibility would be that CLIP-170 foci split off from the leading network in a process resembling fission of condensates.

In this study, we use tandem in-cell and *in vitro* approaches to investigate +TIP condensation and their function. We provide evidence that +TIP networks in cells may have liquid properties. We demonstrate that the +TIPs EB3 and CLIP-170 undergo LLPS at nanomolar concentrations in the absence of crowding agents. The EB3 and CLIP-170-containing droplets co-condense tubulin, and depend on protein regions promoting multivalent interactions. In the presence of microtubules, EB3/CLIP-170 promote rapid growth speeds while reducing catastrophe and pausing frequencies at the microtubule plus-end. Taken together, our results suggest a mechanism whereby +TIP LLPS plays a role in regulating microtubule dynamics.

## Results

### +TIP-networks exhibit properties reminiscent of phase separation

We first studied if +TIP-network profiles with trailing protein foci could result from LLPS, by overexpressing GFP-CLIP-170 in RPE-1 cells. Analysis of plus-end fluorescence intensity profiles in cells revealed that 80 % of the GFP-CLIP-170 networks left behind foci (Figures 1B and C). Dynamic analysis of CLIP-170 networks showed that the remaining protein foci bound to the microtubule shaft dissolved over time (Figure 1D and Movie 1). We next probed the material properties of +TIP-networks by treating cells with the aliphatic alcohol 1,6-hexanediol to reduce hydrophobic interactions (Kroschwald et al., 2017). While untreated cells exhibited 3 µm long +TIP-networks, upon hexanediol treatment, the profile of the +TIP-network reduced to ∼1.5 µm foci (Figures S1A, B and Movie 2), indicating that hydrophobic interactions influence +TIP network organization.

To rule out that the observed foci formation is not an artefact of overexpression, we measured plus-end fluorescence intensity profiles of endogenous CLIP-170. In line with overexpression studies, we observed that for endogenous CLIP-170, ∼70% of the +TIP-networks had foci (Figures 1B and C). EB3, another prominent +TIP, followed a comparable fluorescence profile, with ∼75% of the endogenous EB3 networks having foci (Figures 1B and C), consistent with an overexpression-based study (Mustyatsa et al., 2019). Co-staining of EB3 and CLIP-170 revealed that most networks displayed fluorescence profiles with distinct EB3 and CLIP-170 foci co-localization (Figure 1F).

When we overexpressed the CLIP-170 truncated H2- and H1-mutants (Figure 1A), both tracked the growing GTP-microtubule tip, as previously reported (Figure 1E) (Komarova et al., 2002; Folker et al., 2005). However, the presence of foci behind the leading edge of the H2- and H1-networks was dramatically reduced compared to CLIP-170 (Figure 1C). This suggests that the C-terminal region of CLIP-170 influences network organization at microtubule ends. Since H1 and H2 retain binding to the growing GTP-tip, this result implies that foci formation cannot be solely explained by the presence of GTP-islands in the shaft and might also result from material properties of the multivalent +TIP-network.

### CLIP-170 forms biomolecular condensates in cells

Analysis of +TIP-network fluorescence intensity revealed that with increasing GFP-CLIP-170 expression levels, the CLIP-170 concentration plateaued in +TIP-networks, after an initial increase (Figures 2A and B). This indicates that CLIP-170 in +TIP-networks reaches a state of saturation and suggests a defined stoichiometry of CLIP-170 with the network. Interestingly, at the onset of this transition CLIP-170 patch formed in the cytoplasm (Figures 2A and B). Indeed, overexpression of CLIP-170 in CRISPR/Cas9 knock-in GFP-tubulin RPE-1 cells resulted in cytosolic CLIP-170 patches which displayed liquid properties, undergoing fusion and fission within 5 sec (Figure 2C and Movies 3, 4). Fluorescence recovery after photobleaching (FRAP) revealed that CLIP-170 diffuses highly within these patches and showed a high protein exchange rate with the pool outside the patch, with a recovery half-life and mobile fraction of 27.5 seconds and 93 ± 32 %, respectively (Figure 2D and Movie 5). These properties lead us to conclude that CLIP-170 patches are biomolecular condensates in cells, hereafter called droplets.

**Figure 2:**
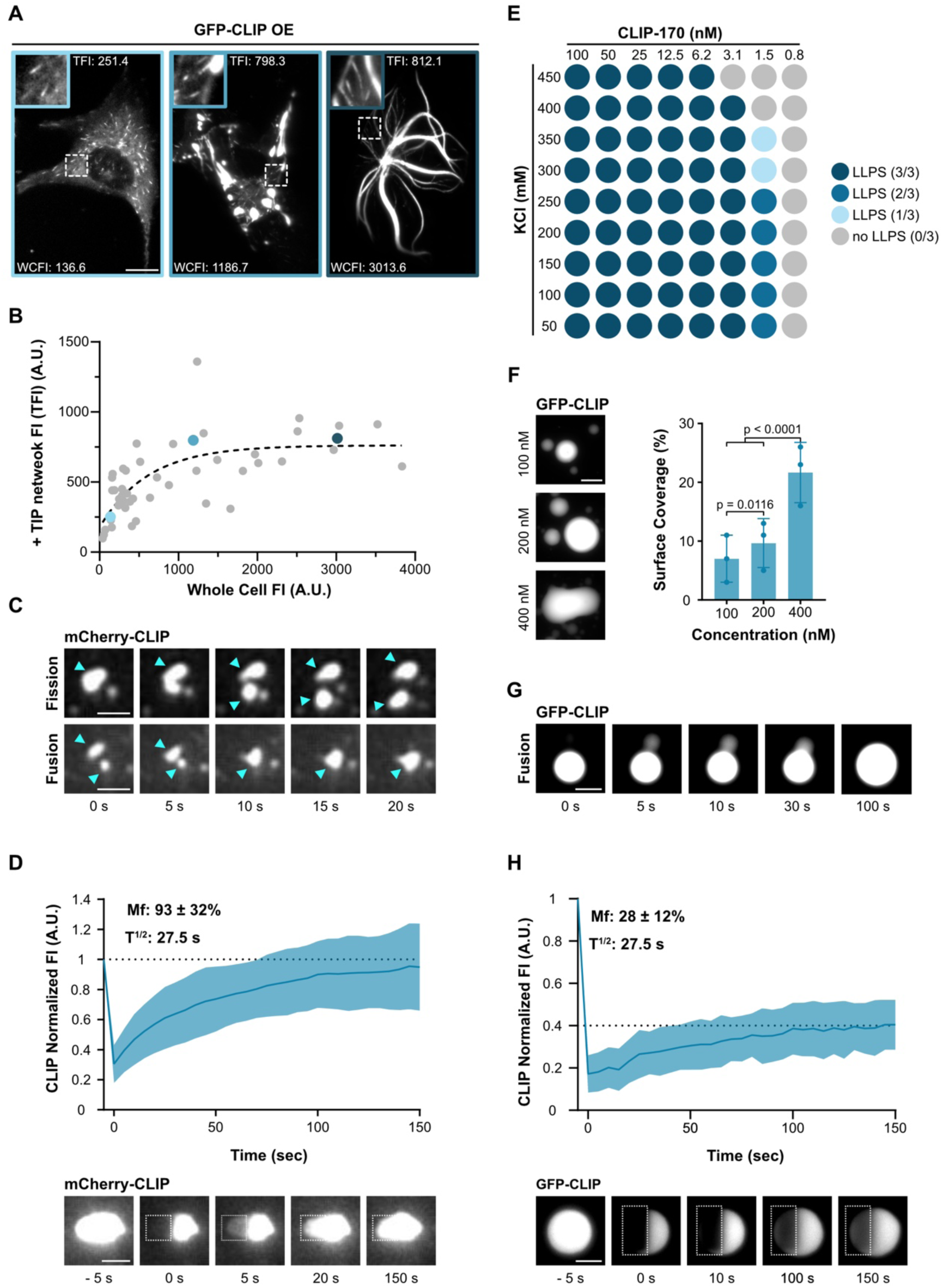
CLIP-170 condenses into droplets in cells and *in vitro*. (**A**) Representative TIRF images of a RPE-1 cells transfected with GFP-CLIP-170, at three different overexpression levels (see also Figure S2A). Zoom-in shows +TIP-networks with adjusted contrast for visualization. For each cell, the whole cell fluorescence intensity (WCFI) and peak of +TIP network fluorescence intensity (TFI) is indicated, blue outlining of the image corresponds to colored data points in (B). Scale bar: 10 µm. (**B**) Analysis of A showing the correlation between peak +TIP network fluorescence intensity and whole cell fluorescence intensity in RPE-1 cells expressing GFP-CLIP-170. Dashed line shows exponential curve fit. Each dot represents 5 analyzed +TIP-networks from one cell, data from 2 independent experiments with a total of 42 cells. (**C**) Representative TIRF time-lapse images of mCherry-CLIP-170 droplets undergoing fission (top panel, cyan arrowheads), and fusion (bottom panel, cyan arrowheads) in cells. Scale bar: 2 µm. (**D**) Representative TIRF images and recovery curve of mCherry-CLIP-170 patches after photobleaching (dashed box). Curve shows mean with SD of 5 individual experiments with a total of 38 droplets from 23 cells. Scale bar: 2 µm. Note that mCherry-CLIP-170 and GFP-CLIP-170 showed the same FRAP-recovery and patch formation behavior (Figures S2A-C) (**E**) Phase diagram of GFP-FL-CLIP *in* vitro at increasing KCl and protein concentration. Blue shaded dot denotes where phase separation occurred, results of 3 independent experiments. (**F**) Representative confocal images of purified GFP-FL-CLIP at indicated concentrations and quantification of the coverslip surface coverage. Statistics: two-tailed Student’s *t*-test. Mean with SD from 3 independent experiments with a total of 27 fields of view per condition. Scale bar: 20 µm. (**G**) Time-lapse images of purified GFP-FL-CLIP (1 µM) undergoing fusion. Representative of 3 experimental replicates. Scale bar: 10 µm. (**H**) Representative images and recovery curve of purified GFP-FL-CLIP (2 µM) droplets after photobleaching (dashed box). Curve shows mean with SD of 3 individual experiments with a total of 47 condensates. Scale bar: 5 µm.

### CLIP-170 forms biomolecular condensates in vitro

To address whether CLIP-170 alone undergoes phase separation, we purified recombinant full-length human GFP-CLIP-170 (FL-CLIP) from insect cells (Figure S3A) and reconstituted its phase separation properties *in vitro*. In the absence of crowding agents and at physiological salt concentrations, FL-CLIP robustly condensed into spheres at concentrations as low as 3.1 nM (Figures 2E and S3B), which is below the cellular concentration of ∼110 nM (Wisniewski et al., 2014). To estimate the amount of FL-CLIP in the dense phase, we used a high throughput confocal microscopy-based approach. Briefly, i) protein mixtures were incubated in 384-well plates, ii) proteins in the dense phase were centrifuged onto the bottom of the wells, iii) wells were imaged using an automated confocal microscope, and iv) surface coverage of protein in the dense phase on well-bottoms was measured (Figure S3C; see Methods for details). A hallmark property of LLPS-potent proteins is that they condense in a concentration-dependent manner and in response to the presence of crowding agents (Alberti et al., 2019). In line with this, the sphere size and amount of FL-CLIP in the dense phase increased in a concentration-dependent manner even in the absence of crowding agents (Figures 2F and S3D). Increasing the ionic strength of the buffer reduced the sphere size (Figure S3B). Addition of a crowding agent, 2 % polyethylene glycol (PEG), increased the amount of FL-CLIP in the dense phase by 1.5-fold but reduced sphere size (Figures S3E and F).

In line with the case in cells, FL-CLIP spheres displayed liquid properties such as fusion, although 5-times slower than what we observed in cells (Figure 2G and Movie 6). We further interrogated the liquid properties of these spheres by FRAP, which confirmed that FL-CLIP diffuses within the spheres with a half-life of 27.5 s, identical to the recovery speed in cells. However, FL-CLIP exchange dynamics were reduced three-fold with a mobile fraction of 28 ± 12% (Figure 2H and Movie 7). By calibrating the fluorescent intensity of GFP, we estimated that the initial FL-CLIP concentration increased 8.5-fold within these spheres (Table 1). Collectively, these results show that FL-CLIP undergoes LLPS and forms droplets *in vitro* at nanomolar concentration and in the absence of further proteins or crowding agents. We hypothesize that the discrepancies between the material properties of the droplets in cells and *in vitro* are due to the presence of additional proteins and/or CLIP-170 post-translational modifications in the cell droplets.

**Table 1:** CLIP-170, EB3 and tubulin concentration inside droplets. Protein concentration outside droplets (C_dilute_) and inside droplets (C_droplet_), obtained by calibrating fluorescent intensities of GFP, mCherry and Atto-565 see calibration in Figure S9.

| | $C_{\text{dilute}}$ (nM) | $C_{\text{droplet}}$ (nM) | Ratio ($C_{\text{droplet}}/C_{\text{dilute}}$) |
| --- | --- | --- | --- |
| **CLIP-170** ( 200 nM) | 70 | 1 700 | 24 |
| **mcherry-EB3** (2 μM) + EB3 (8 μM) | 8 600 | 31 000 | 3.5 |
| **CLIP-170** (200 nM) + mcherry-EB3 (1 μM) | 75 | 1 800 | 24 |
| **mcherry-EB3** (1 μM) + CLIP-170 ( 200 nM) | 400 | 2 000 | 5 |
| **CLIP-170** (200 nM) + tubulin (400 nM) | 50 | 1 700 | 34 |
| **Tubulin** (400 nM) + CLIP-170 (200 nM) | 300 | Droplet 650 / Shell 1 200 | Droplet 2 / Shell 4 |

### The CLIP-170 C-terminal region drives CLIP-170 into the dense phase

We next investigated which domains of CLIP-170 drive droplet formation. We expressed GFP-tagged H2- and H1-CLIP mutants in RPE-1 cells where the C-terminal region is truncated or fully removed (Figure 1A) (Pierre et al., 1992, 1994; Diamantopoulos et al., 1999; Goodson et al., 2003). In line with previous observations, we saw that H1 and H2 displayed microtubule plus-end tracking activity in cells, but did not form any cytosolic droplets even at high expression levels (Figure 3A and Movies 8, 9) (Pierre et al., 1994; Goodson et al., 2003). To understand to which extent the C-terminal region is necessary for CLIP-170 to undergo LLPS, we purified recombinant human H1 and H2 from bacteria and measured their ability to condense *in vitro* at nanomolar concentrations (Figure S4A). While FL-CLIP phase separated in the absence of other factors, H1 showed faint irregular-shaped aggregation but no droplet formation, even in presence of 10 % PEG (Figures 3B, C and S4B, C). In contrast to the case in cells, at nanomolar concentration H2 underwent condensation *in vitro*, although with a 300-fold reduction of protein surface coverage and a 25-fold reduction in droplet size compared to FL-CLIP (Figures 3B, C, D and S4A). Addition of 2 % PEG increased H2 condensate formation by 3-fold but did not affect condensate size (Figures 3B, C and S4B, C).

**Figure 3:**
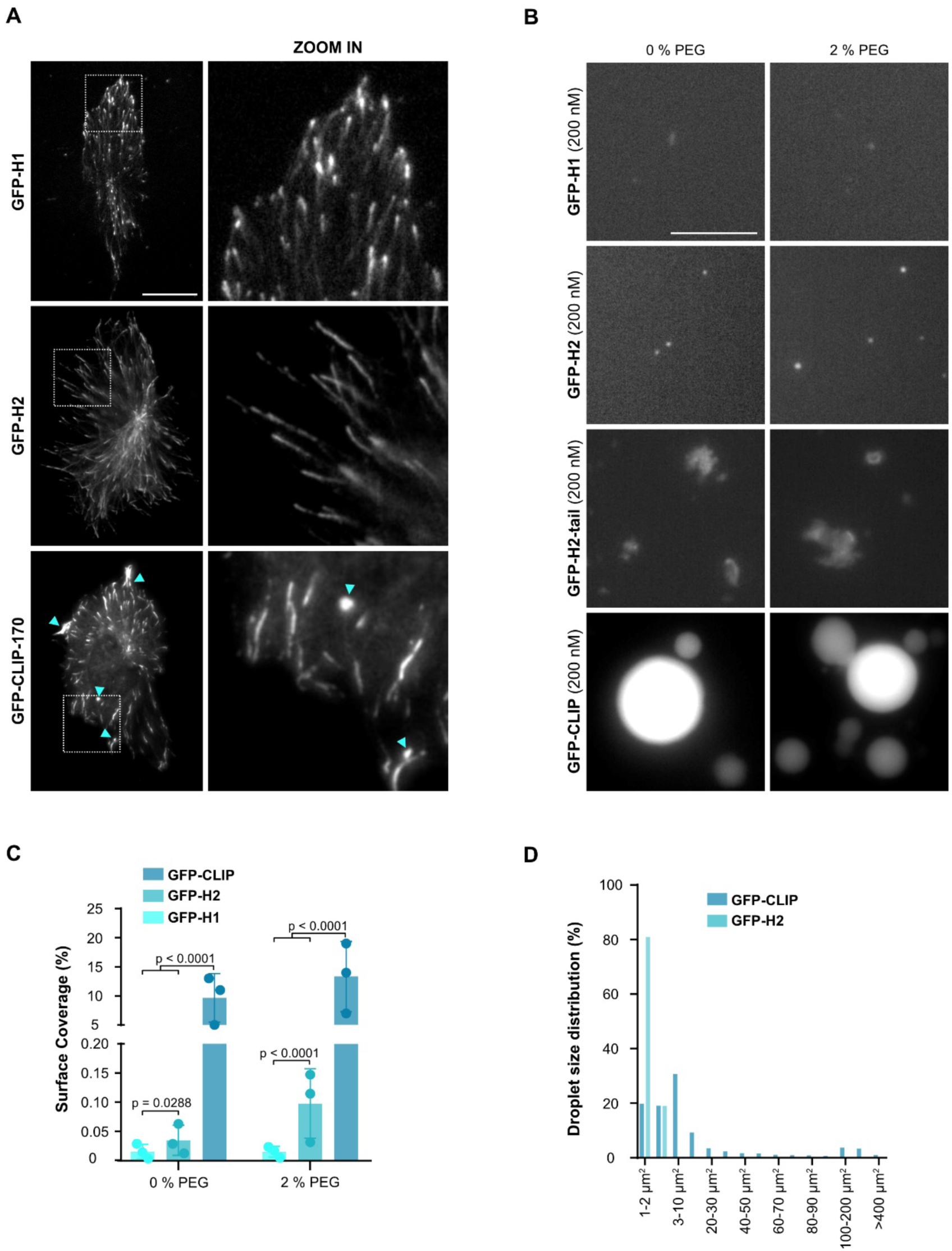
The C-terminal region drives CLIP-170 into the dense phase. (**A**) Representative images of fixed RPE-1 cells transfected with full length GFP-CLIP-170, GFP-H2 or GFP-H1 (left panel) with insets (right panel). Cyan arrowheads denote droplet formation in GFP-CLIP-170 expressing cell. Scale bar: 20 µm. (**B**) Representative confocal images of purified GFP-H1, GFP-H2, GFP-H2-tail and GFP-CLIP-170 each at 200 nM in the absence (left panel) or presence (right panel) of 2 % PEG. Scale bar: 20 µm. **(C)** Condensate surface coverage of the three constructs at indicated PEG concentrations. Mean with SD from 3 independent experiments with a total of 27 fields of view per condition. Statistics: two-tailed Student’s *t*-test. (**D**) Size distribution of GFP-FL-CLIP (200 nM) and GFP-H2 (200 nM) droplets in the absence of PEG. Graph shows average size distribution from 3 independent experiments with a total of 27 fields of view.

Intrinsically disordered domains are common features of proteins that undergo LLPS (Boeynaems et al., 2018). To parse out the contributions of disordered domains to FL-CLIP’s phase separation potency, we engineered a mutant containing the H2 domain fused to the far C-terminal region, termed H2-tail (Figure S4E). This mutant lacks the majority of the coiled-coil region but retains many predicted disordered regions within CLIP-170 (Figure S4E). Attempts to purify this mutant resulted in insoluble large protein aggregates, highlighting the importance of the coiled-coil region for CLIP-170’s solubility (Figure 3B). Collectively, these results show that while the monomeric H1 is sufficient to track the growing microtubule tip, the dimeric form of CLIP-170 is necessary to undergo LLPS and that the C-terminal region robustly drives CLIP-170 condensation.

### EB3 undergoes LLPS and co-condenses with CLIP-170

CLIP-170 requires the presence of EBs to localize to growing microtubule ends (Dixit et al., 2009, Bieling et al., 2008). This prompted us to ask whether purified FL-CLIP and EB3 could co-condense *in vitro*. We first studied EB3 alone and observed that it has the capacity to undergo LLPS at micromolar concentrations: compared to 48 ± 17 µm^2^ large droplets observed for 400 nM FL-CLIP, at 1 µM EB3 phase separated into many small 1.9 ± 0.6 µm^2^ droplets, resulting in a 3.5-fold reduced surface coverage (Figures 4A, B and S5A). EB3 fraction in the dense phase was comparable between 0 and 2 % PEG (Figures 4A and B). FRAP revealed that EB3 diffused within these patches at the same rate as FL-CLIP, but had a 2.5-fold higher protein exchange rate with the pool outside the droplet (Figure 4C and Movie 10).

**Figure 4:**
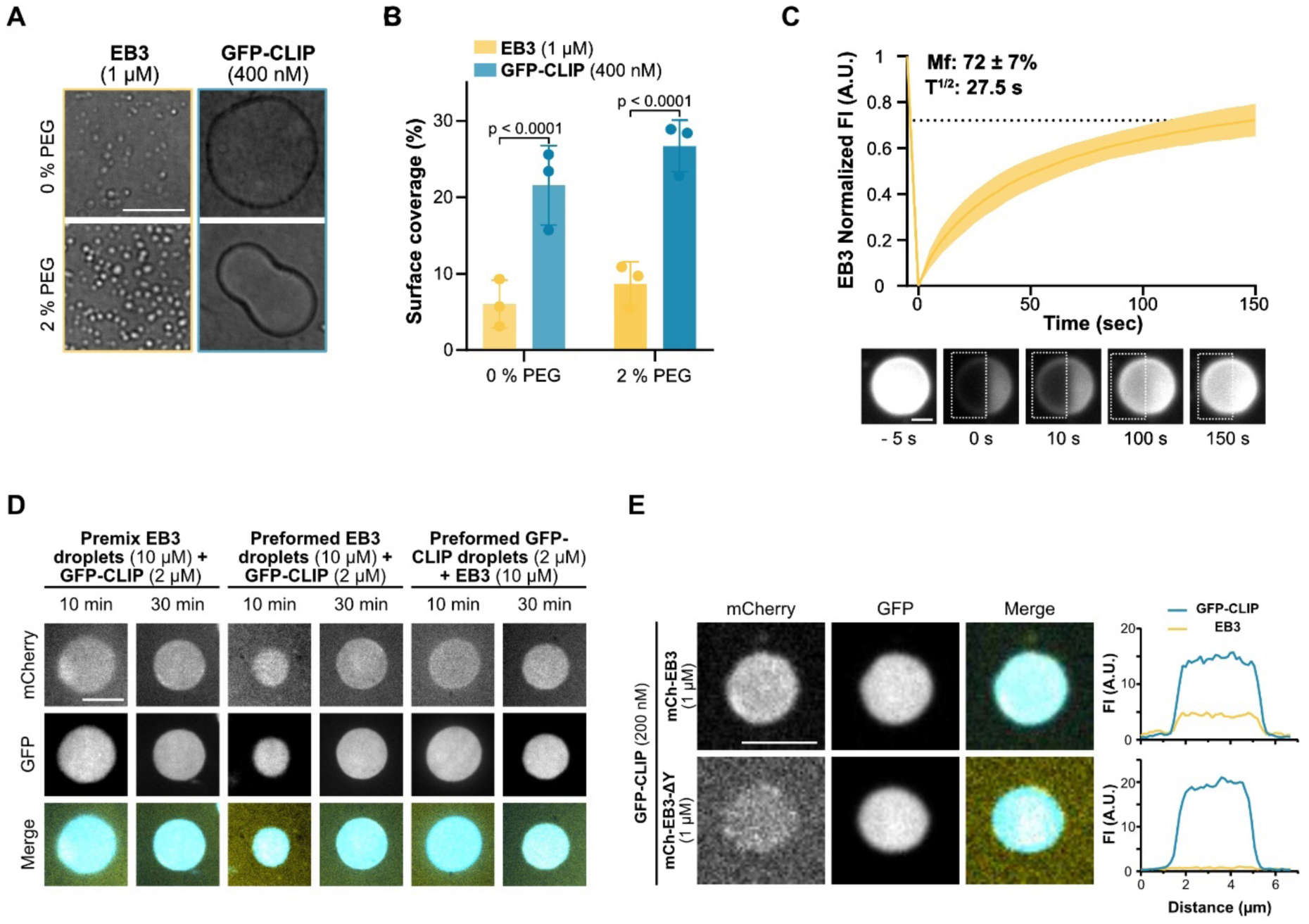
EB3 undergoes LLPS and co-condenses with CLIP-170 *in vitro*. (**A**) Representative DIC images of purified EB3 (1 µM) and GFP-FL-CLIP (400 nM) in absence (top panel) or presence (bottom panel) of 2 % PEG. Scale bar: 20 µm. (**B**) Condensate surface coverage of EB3 (1 µM) and GFP-FL-CLIP (400 nM) at indicated PEG concentrations. Mean with SD from 3 independent experiments with a total of 27 fields of view. Statistics: two-tailed Student’s *t*-test. (**C**) Representative images and recovery curve of purified EB3 (10 µM) + mCherry-EB3 (100 nM) droplets after photobleaching (dashed box). Curve shows mean with SD of 3 independent experiments with a total of 32 condensates. Scale bar: 5 µm. (**D**) Representative images of 2 µM GFP-FL-CLIP and 10 µM EB3 (8 µM unlabeled + 2 µM mCherry-EB3) in the denoted mixing conditions; 10 min after mixing and at coarsy steady state - 30 min after mixing. Scale bar: 10 µm. (**E**) Representative images of EB3/GFP-FL-CLIP droplets (top) or EB3-Λ1Y/GFP-FL-CLIP droplets (bottom) at denoted concentrations with the corresponding line scan. Scale bar: 4 µm.

We next tested the ability of CLIP-170 to co-condense with EB3. Indeed, FL-CLIP and EB3 robustly co-condensed into droplets, in which both proteins showed a homogenous distribution (Figure 4D and S5B). To address if diffusion kinetics are impacted by a multivalent EB3/FL-CLIP-network, we repeated our FRAP experiment. Surprisingly, the time to recover after photobleaching was the same for EB3, FL-CLIP and EB3/FL-CLIP droplets (Movies 7, 10, 11 and Figures 2H, 4C and S5C). The mobile fraction of FL-CLIP and EB3/FL-CLIP was also comparable, however EB3 droplets had a 2-fold higher mobile fraction. This indicates that fluorescent recovery of FL-CLIP and EB3/FL-CLIP droplets depends more on protein diffusion, while protein exchange with the soluble pool further impacts recovery of EB3 droplets. In line with this observation, the three different droplets exhibited distinct surface interaction properties: EB3 did wet the coverslip resulting in flat droplets, while FL-CLIP and EB3/FL-CLIP droplets remained a similar sphere-like shape when bound to the coverslip (Movie 12, 13 and 14). Co-condensation of EB3 with FL-CLIP increased the number of small droplets, such that droplets larger than 40 µm^2^ were rarely observed (Figures S5D). This reduced the average droplet size by 20-fold compared to FL-CLIP, despite a 6-fold increase in total protein concentration in the solution (2 µM to 12 µM) (Figure S5E). These results show that EB3 and FL-CLIP can co-condensate and that the multivalency of the network does not further impact FL-CLIP diffusion within the network, while droplet size and surface tension can be impacted by the protein composition.

To study whether partitioning of +TIPs into pre-formed droplets affected condensation properties, we compared: i) co-condensation of EB3 and FL-CLIP, ii)) addition of FL-CLIP to pre-formed EB3 droplets and iii) addition of EB3 to pre-formed FL-CLIP droplets. Addition of FL-CLIP to EB3 droplets increased the droplet size over time; however, upon reaching coarsy steady state, we observed no differences in droplet size or protein concentrations within the droplets between the three conditions (Figure 4D).

Interactions between EB1 and CLIP-170 are driven by the last C-terminal tyrosine of EB1 (Bieling et al., 2008). To probe whether specific protein-protein interactions underlie EB3/FL-CLIP co-condensation, we purified a EB3 tyrosine deletion mutant (EB3-ΔY) and repeated our colocalization assay. Removal of the last tyrosine drastically reduced EB3’s ability to co-condense with FL-CLIP (Figure 4E), despite leaving EB3-ΔY’s intrinsic phase separation properties intact (Figure S5F). Thus, specific binding interactions between EB3 and FL-CLIP are key features in their ability to co-condense. Altogether, these data show that EB3 undergoes phase separation and can co-condense with FL-CLIP into droplets with homogenous protein distribution. These droplets exhibit similar fluid properties, but likely different surface interaction properties.

### EB3/CLIP-170-networks cooperatively phase separate

Given that EB3/FL-CLIP droplets were reduced in size, we quantified whether the total amount of protein in the dense phase is reduced. Measuring the concentration of FL-CLIP in droplets compared to the dilute phase showed that FL-CLIP is enriched 24-fold in the droplet (Table 1). Co-condensation of FL-CLIP and EB3 only slightly increased the concentration of FL-CLIP in the droplet, while condensation of EB3 was increased 1.4-fold (Table 1). This implies, that FL-CLIP drives the partitioning of EB3 into droplets. To probe this observation further and to analyze the total amount of protein in the diluted and dense phase, we performed a droplet-pelleting assay followed by SDS-PAGE analysis (for details see Methods). Similar to what we observed by microscopy, in EB3/FL-CLIP networks the amount of condensed EB3 increased by 1.7-fold compared to amounts measured for EB3 alone (Figures 5A and B). We further studied the synergistic effect on droplet formation with our high-throughput microscopy assay. Although the size of EB3/FL-CLIP droplets decreased, the number of droplets increased leading to a surface coverage 2.5-fold higher than FL-CLIP droplets alone (Figure 5C, D and S5E). This resulted in a 40 % increase of surface coverage by EB3/FL-CLIP networks when compared to the sum of the surface coverages of EB3 alone plus FL-CLIP alone (Figure 5E).

**Figure 5:**
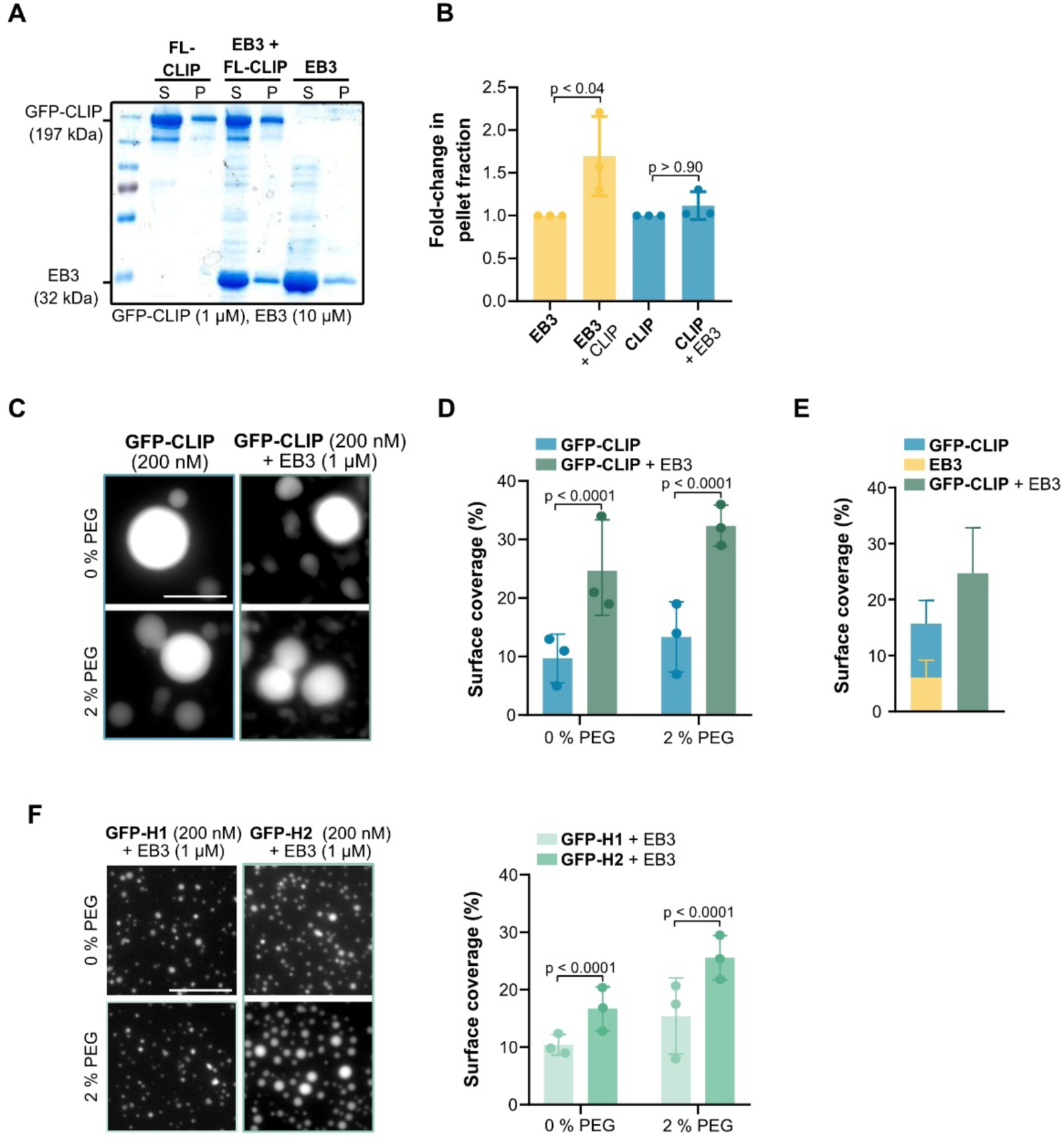
Synergistic condensation of CLIP-170 and EB3. (**A**) Representative SDS-PAGE analysis from the droplet-pelleting assay showing protein fractions in supernatant dilute phase (S) or pellet dense phase (P) under each condition: 1 µM GFP-FL-CLIP; 10 µM EB3; EB3/GFP-FL-CLIP (10 µM + 1 µM). (**B**) Quantification of SDS-PAGE analysis showing the fold-change of protein in the pellet fraction at the three conditions. Mean with SD from three independent experiments. Statistics: one-way ANOVA. (**C**) Representative fluorescence confocal images of purified GFP-FL-CLIP in the absence (left) or presence (right) of EB3, and in the absence (top panel) or presence (bottom panel) of 2 % PEG. Scale bar: 20 µm. (**D**) Condensate surface coverage of purified GFP-FL-CLIP in the absence (left) or presence (right) of EB3 at indicated PEG concentrations. Mean with SD from 3 independent experiments with a total of 27 fields of view. Statistics: two-tailed Student’s *t*-test. (**E**) Quantification of droplet surface coverage of EB3 and GFP-FL-CLIP alone compared to surface coverage of EB3/GFP-FL-CLIP droplet formation when undergoing synergistic LLPS in the absence of PEG. (**F**) Representative fluorescence confocal images and quantification of purified GFP-H1 (left) or GFP-H2 (right) in the presence of EB3, and in the absence (top panel) or presence (bottom panel) of 2 % PEG. Scale bar: 20 µm. Right: condensate surface coverage of indicated GFP-H1 (200 nM) or GFP-H2 (200 nM) in the presence of EB3 (1 µM), and in the presence of the indicated PEG concentrations. Mean with SD from 3 independent experiments with a total of 27 fields of view. Statistics: two-tailed Student’s *t*-test.

Repeating the EB3 experiment with H2 reduced surface coverage by 1.5-fold compared to the presence of FL-CLIP, and H1 further decreased surface coverage by 2.5-fold (Figures 5F and S5G). These data demonstrate that CLIP’s C-terminal region is essential for a highly multivalent EB3/FL-CLIP network formation. We further show that EB3 and CLIP-170 can undergo LLPS both independently, and that when acting as an ensemble, the amount of proteins in the dense phase synergistically increases.

### CLIP-170 and EB3 condense tubulin in vitro

A CLIP-170 dimer has as many as 8 tubulin binding sites, can bind tubulin *in vitro,* and colocalizes with tubulin in cells (Figure 6A) (Pierre et al., 1994 et al., 1999; Perez et al., 1999; Gupta et al., 2010). We therefore asked if CLIP-170 droplets can co-condense tubulin *in vitro*. Tubulin alone did not form droplets at micromolar concentrations; even in the presence of 5% PEG only aggregation was observed (Figure S6A). However, when 200 nM FL-CLIP was mixed with 400 nM Atto565-tubulin, the two proteins phase separated into 26.8 ± 12 µm^2^ droplets (Figures 6B and S6B). We repeated these experiments with H1 and found irregular-shaped aggregation with tubulin but no droplet formation (Figure 6B). Based on the intrinsic phase separating capacity of EB3, we studied whether EB3 alone could condense tubulin (Figure 4A). Indeed, EB3 also co-condensed tubulin, although into much smaller 1 ± 0.07 µm^2^ droplets, in line with our observed EB3 droplet size distribution (Figures 6C and S6C). EB3/FL-CLIP networks also condensed tubulin into intermediate-sized droplets, consistent with EB3 reducing the droplet size compared to FL-CLIP (Figure S6C). These results demonstrate that FL-CLIP, EB3 and EB3/FL-CLIP networks are potent to condense tubulin.

**Figure 6:**
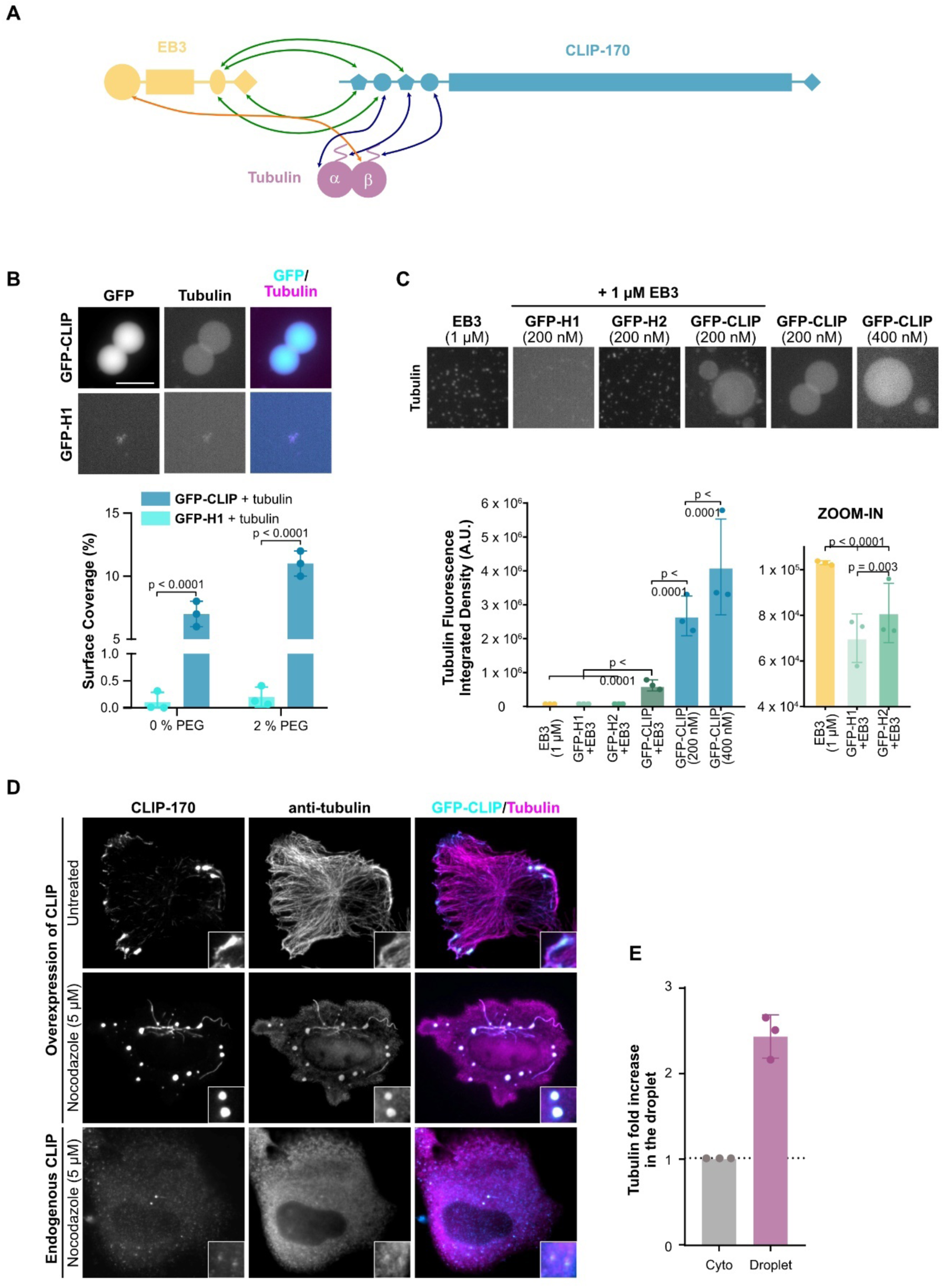
CLIP-170 and EB3 droplets condense tubulin. (**A**) Cartoon schematic of domain interactions between EB3, CLIP-170, and tubulin based primarily on (Gupta et al., 2010; Chen et al., 2021; Bjelic et al., 2012). Interaction sites between EB3 and CLIP-170, EB3 and tubulin, and CLIP-170 and tubulin are shown with green, orange, and blue arrows, respectively. For simplification, monomers of EB3 and CLIP-170 are shown. (**B**) Top: representative confocal images of purified GFP-FL-CLIP and GFP-H1 each at 200 nM with Atto-565-tubulin (400 nM). Scale bar: 20 µm. Bottom: quantification of the coverslip surface coverage of tubulin in presence of GFP-FL-CLIP or GFP-H1. Mean with SD from 3 independent experiments with a total of 27 fields of view. Statistics: two-tailed Student’s *t*-test. Note that co-condensation is independent of the fluorescent tag, as unlabeled tubulin also partitioned into GFP-FL-CLIP droplets see Figure S6B. (**C**) Top: representative confocal images of Atto-565-tubulin (400 nM) in the presence of purified EB3 (1 µM), GFP-H1 (200 nM) and EB3 (1 µM), GFP-H2 (200 nM) and EB3 (1 µM), GFP-FL-CLIP (200 nM) and EB3 (1 µM), and GFP-FL-CLIP (200 nM and 400 nM) alone. Scale bar: 20 µm. Bottom: quantification of the integrated density of tubulin fluorescence under denoted conditions with zoom in for the first three conditions. Mean with SD from 3 independent experiments with a total of 27 fields of view. Statistics: two-tailed Student’s *t*-test. (**B-C**) Note that droplet centrifugation onto the coverslip lead to a homogenous distribution of tubulin in the GFP-FL-CLIP droplet, see Figure S6B (**D**) Representative images of fixed RPE-1 cells transfected with GFP-CLIP-170 and untreated (top panel) or treated with 5 µM nocodazole for 1 hour (middle panel); RPE-1 WT cells were treated with 5 µM nocodazole (bottom panel) and stained for endogenous CLIP-170 and tubulin. Scale bar: 10 µm. Images are representative of 3 independent experiments. **(E)** Graph showing normalized tubulin fluorescence intensity in CLIP-170 droplets compared to cytoplasm in full-length GFP-CLIP-170 transfected RPE-1 cells treated with nocodazole. Mean with SD from 3 independent experiments with a total of 126 condensates from 26 cells.

Although all three different protein droplets condensed tubulin, the distribution of tubulin differed. Tubulin was homogenously mixed within FL-CLIP droplets and in addition formed a shell-like structure around the droplets (Figure S6D). Within the shell, tubulin was concentrated twice as high as within the droplet (Table 1). This result indicates that within the droplet tubulin is mixed with FL-CLIP, while tubulin phase separates at the solution interface. In contrast to FL-CLIP droplets, tubulin was homogenously distributed in EB3 and EB3/FL-CLIP droplets (Figure S6D). This implies that addition of EB3 changes the surface properties such that tubulin condensation at the interphase is not favorable.

Strikingly, FL-CLIP droplets enriched condensed tubulin 40-fold more than EB3 droplets, even at 5-fold lower FL-CLIP concentrations (Figure 6C). This difference in condensed tubulin cannot be solely explained by the 3.5-fold difference in surface coverage between EB3 and FL-CLIP, but indicates a different in the potency of the two proteins to condense tubulin (Figure 6C). Based on this result, we asked whether addition of EB3 to FL-CLIP impacts tubulin condensation. Indeed, we observed that EB3/FL-CLIP networks condensed tubulin 4-fold less efficient than FL-CLIP alone (Figure 6C). We hypothesize that within a multivalent tubulin/EB3/FL-CLIP network the heterotypic interaction between EB3 and FL-CLIP are stronger than EB3-tubulin and FL-CLIP-tubulin interactions, leading to a reduction of tubulin condensation, especially around the droplets. Collectively, these results show that FL-CLIP can condense tubulin effectively and that EB3 reduces the tubulin condensation capacity of FL-CLIP droplets, while changing the surface properties of the droplets.

### Tubulin co-condenses with CLIP-170 in cells

We next addressed whether CLIP-170 can condense tubulin in cells. Overexpressing GFP-CLIP-170 in RPE-1 cells revealed that CLIP-170 droplets colocalized with areas of high tubulin fluorescence intensity (Figure 6D, top panel) (Perez et al., 1999). However, it was not possible to distinguish if these areas corresponded to microtubule bundles, or if the local increased signal resulted from tubulin condensation. To address this question, we depolymerized the microtubule network with 5 µM nocodazole. After microtubule depolymerization, tubulin showed robust co-localization with CLIP-170 droplets (Figure 6D, middle panel), and tubulin fluorescence intensity was 2.4-fold higher in the droplets compared to the cytoplasm (Figure 6E). To study whether CLIP-170 could also co-condense tubulin at endogenous concentration, we used antibody staining after microtubule network depolymerization. In WT cells, small foci of endogenous CLIP-170 were frequently observed after microtubule network depolymerization, and these foci showed local enrichment of tubulin (Figure 6D, bottom panel). In cells depleted of CLIP-170 and in CLIP-170 knockdown cells rescued with H2, we did not observe tubulin/FL-CLIP foci (Figure S6E). Contrary to our *in vitro* experiments, tubulin foci were not observed in nocodazole-treated cells overexpressing H2 or EB3 (Figure S6F). After depolymerization of the microtubule network EB3 was cytoplasmic with a few small foci, but showed no distinct co-condensation with tubulin (Figure S6F). We conclude that CLIP-170, but not EB3, can condense tubulin in cells.

### Phase separation-potent +TIP networks increase microtubule growth

The absence of either EB3 or CLIP-170 at the microtubule tip does not reduce growth speeds in cells (Komorova et al., 2002; Straube et al., 2007; Komorova et al., 2009). However, combined siRNA knockdown of EB3 and CLIP-170 reduced microtubule growth speeds by 20 % (Figures S7C and D). To understand if LLPS of +TIPs could impact microtubule tip dynamics, we turned to *in vitro* reconstitution. We first reconstituted microtubule growth in presence of +TIP-networks with either reduced (50 nM H2 + 800 nM EB3) or minimal (50 nM H1 + 800 nM EB3) LLPS activity (Figure 5F), implying reduced or minimal co-condensation of tubulin at the growing tip (Figure 6C). To our knowledge these CLIP mutants preserve all binding domains to interact with EB3 and tubulin, implying that the difference between these networks lies primarily in their phase separation potency. As observed previously, EB3 alone increased microtubule growth speeds by 1.5-fold and catastrophe frequency by 8-fold (Figures 7A and B) (Komorova et al., 2009; Montenegro Gouveia et al., 2010). Addition of H1 or H2 to the assay with EB3 resulted in tip-tracking behavior by EB3/H1 and EB3/H2 networks, but did not further change any of the dynamic parameters compared to EB3 alone (Figures 7A, B and S8A). These results show that CLIP/EB3 networks with reduced or minimal capacity to condense tubulin have the same impact on microtubule tip dynamics as EB3 alone.

**Figure 7:**
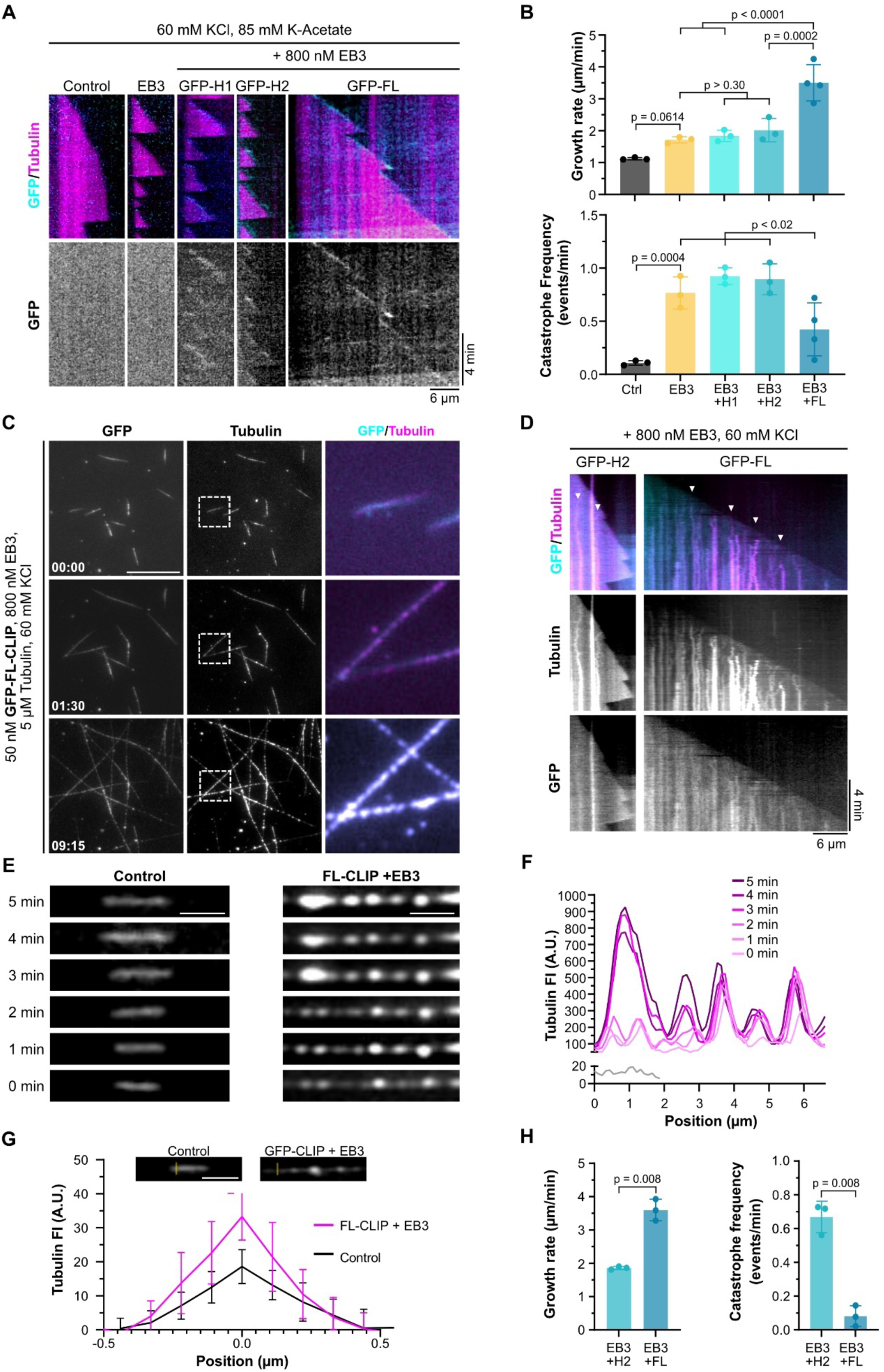
LLPS of +TIPs regulates microtubule dynamics through local tubulin condensation. (**A**)Representative microtubule kymographs of denoted +TIP-networks in higher salt buffer (60 mM KCl and 85 mM K-acetate, see materials and methods). Note that tip-tracking efficiency (GFP channel) is weaker at 5 µM tubulin than at higher tubulin concentrations (Figure S8B). (**B**) Microtubule growth rate (top) and catastrophe frequency (bottom) in presence of denoted proteins in high salt buffer. Mean with SD of minimum of three independent experiments with the following number of analyzed microtubules: Control – 48; EB3 – 31; EB3/H1 – 28; EB3/H2 – 28; EB3/FL-CLIP - 60. Statistics: One-way ANOVA Fisher’s LSD test. (**C**) Representative time-lapse TIRF images of GFP-FL-CLIP (50 nM) with Atto-565-tubulin (5 µM) in the presence of unlabeled EB3 (800 nM) and 60 mM KCl. Time denoted in minutes: seconds; scale bar: 20 µm. The zoom-ins (white dashed box) show representative droplet formation along microtubules. (**D**) Representative microtubule kymographs in the presence of GFP-H2 or GFP-FL-CLIP (50 nM) and EB3 (800 nM) grown at 5 µM tubulin and 60 mM KCl. Arrowheads denote areas of robust tubulin/FL-CLIP condensation on growing microtubule shaft. (**E**) Representative images of Atto-565-tubulin microtubules growing in the absence (left, control) and in presence of EB3/FL-CLIP (right). Right images show tubulin condensation along microtubule over time. Scale bar: 2 µm. (**F**) Corresponding line scan of E with the gray line scan representing the 5 µM tubulin control condition and the magenta line scans of 5 µM tubulin in presence of EB3/FL-CLIP (800 nM/50 nM). (**G**) Tubulin fluorescence intensity in tip-proximal regions in 5 µM tubulin control condition and in presence of EB3/FL-CLIP. Quantification of perpendicular line scans in tip-proximal regions. Mean with SD from 2 independent experiments for each condition and a total of 20 microtubules; Scale bar: 2 µm. (**H**) Quantification of microtubule growth rate (left) and catastrophe frequency (right) in the presence of EB3/H2-networks and EB3/FL-CLIP droplets in experiments from D (60 mM KCl). Mean with SD of three independent experiments with the following number of analyzed microtubules: EB3/H2 – 29; EB3/FL-CLIP – 59. Statistics: paired t-test.

When we repeated the reconstitution experiments with EB3/FL-CLIP, microtubules grew at a speed of 3.5 µm/min, a 1.5-fold increase compared to analogous experiments with EB3/H2 and EB3/H1, and nearly four-fold increase compared to controls with tubulin alone (Figures 7A, B and Movie 15). Furthermore, catastrophe events were reduced under these conditions (Figures 7A and B), and when they occurred were rapidly followed by rescue events (Figure S8A). To reconstitute this fast microtubule growth in presence of a phase separation deficient EB3/H2 network, we increased the tubulin concentration while keeping the EB3/H2 concentration constant. We measured microtubule growth speeds and found that a growth speed of 3.5 µm/min (the speed achieved by EB3/FL-CLIP in the presence of 5 µM tubulin) corresponds to 12.8 µM tubulin in presence of EB3/H2 (Figures S8B and C). These results show that phase separation-potent EB3/FL-CLIP networks increase the growth rate and reduce the catastrophe events compared to EB3/H2 networks. A phase separated +TIP network might impact the growing microtubule tip due to changes in material properties, or due to condensation of tubulin at the tip, based on our observation that EB3/FL-CLIP can condense tubulin (Figure 6C).

### +TIPs undergo LLPS and condense tubulin on microtubules, promoting microtubule growth

To understand if LLPS of EB3/FL-CLIP networks could enrich tubulin on microtubules, we used microtubules as a platform to induce LLPS. A recent study showed that addition of cell lysate containing overexpressed CLIP-170 to microtubules formed CLIP-170-containing droplets along microtubules (Jijumon et al., 2022). We asked if this droplet formation is due to the crowding environment of the cell lysate, or a general property of FL-CLIP. Under assay conditions with lower ionic strength (see materials and methods), we repeated our assay from above to study if purified EB3 together with FL-CLIP could phase separate along microtubules. When we incubated purified CLIP-170 and EB3 with dynamic microtubules in vitro, we found that purified EB3/FL-CLIP formed droplets along the shaft (Figure 7C and Movie 16).

In line with our observations that EB3/FL-CLIP droplets condensed tubulin (Figure S6D), these networks enriched tubulin all along the microtubule and co-condensed tubulin over time into droplets on the shaft, even more efficient than free floating droplets (Figures 7C-G). At this condition where tubulin condensed along the microtubule, we observed rapid microtubule growth speeds of 3.6 µm/min and very few catastrophe events or pauses in the growth phase (Figures 7D, H and S8D). This increase in growth speed and reduction in catastrophe events is in line with our results from above where EB3/FL-CLIP was located at the microtubule tip. When we repeated these experiments with less potent phase separating EB3/H2-networks, we only occasionally observed condensate formation along the microtubule (Figure 7D, left panel). In presence of EB3/H2-networks growth speeds were 2-fold slower than for EB3/FL-CLIP, catastrophe frequencies were increased 7-fold and pauses increased 30-fold (Figure 7H and S8D). In the absence of EB3, we did not observe microtubule binding or increased growth speeds for any CLIP constructs (Figures S8E and F). Collectively, these experiments demonstrate that EB3/FL-CLIP networks undergo LLPS on microtubules, can condense tubulin, and drive microtubule growth.

## Discussion

Rearrangements of the microtubule network architecture require spatiotemporal regulation of microtubule growth. A large body of work has highlighted the role of +TIPs that act as microtubule polymerases (such as XMAP215) or increase microtubule tip dynamics (such as the EBs) in promoting these highly regulated changes (Gard and Kirschner, 1987; Srayko et al., 2005; Straube and Merdes, 2007; Brouhard et al., 2008; Bieling et al., 2008; Vitre et al., 2008; Zanic et al., 2013; Yang et al., 2017). While these studies showed the impact of +TIPs on microtubule dynamics, it has remained unclear how these proteins self-assemble to form highly dynamic and multivalent networks. LLPS allows for network formation of proteins based on highly multivalent interactions. We now provide evidence that LLPS of +TIPs could be a mechanism to explain the formation of +TIP-networks at the growing microtubule tip. This mechanism could be a base to locally recruit and enrich diverse +TIPs.

LLPS of proteins is often driven through weak multivalent interactions and low complexity domains (Boeynaems et al., 2018). A growing body of evidence also implicates coiled-coil regions of proteins in driving their LLPS. Coiled-coil domains allow for rapid dimerization and self-oligomerization which might be sufficient for the assembly of certain condensates (Larson et al., 2017; Strom et al., 2017). Notably, coiled-coil containing proteins are highly enriched in phase-separated centrosomes and P-granules (Salisbury, 2003; Ford and Fioriti, 2020), and many of these proteins phase separate in a coiled-coil dependent manner (Mitria and Kriwacki, 2016; Woodruff et al., 2017; Jiang et al., 2021). Our experiments using CLIP-170 truncation mutants emphasize the role that coiled-coil proteins can play in driving LLPS. In addition, our data implies that specific binding domains between EB3 and FL-CLIP are needed to form a multivalent network, as deletion of one single tyrosine strongly reduces EB3 co-condensation with FL-CLIP.

Here, we propose a new mechanism by which LLPS of +TIPs regulate microtubule growth. Our model of LLPS-driven microtubule growth is based on the following observations: i) the +TIPs EB3 and CLIP-170 have the capacity to undergo LLPS *in vitro* and in cells (Figures 2-5), ii) phase separated EB3/FL-CLIP droplets can condense tubulin (Figure 6), iii) microtubule growth speed is increased and catastrophe frequency reduced when EB3/FL-CLIP undergo phase separation on microtubules (Figure 7H), and iv) tubulin condensed along the microtubule in presence of phase separated EB3/FL-CLIP (Figure 7C-G). We hypothesize that this tubulin enrichment may be a mechanism to increase local tubulin availability at microtubule ends uncoupled from cytoplasmic tubulin concentrations. Microtubule dynamics might be further modulated by the viscosity of the droplets, such as sterically constraining protofilaments in a manner that accelerates growth or prevents catastrophes.

Microtubules could serve as a general platform to concentrate proteins locally to a LLPS-sufficient concentration. We hypothesize that the mechanism of LLPS initiation at the growing tip depends on conformational properties of the growing microtubule tip, which leads to differential binding of EB3 to the tip over the shaft (Zhang et al., 2015) and subsequent recruitment of CLIP-170 to form a multivalent network. As LLPS is a concentration dependent process, the local recruitment of EB3 and CLIP-170 could increase their concentration compared to the solution to such an extent that they undergo LLPS. We hypothesize that, upon reaching a critical CLIP-170 concentration, the +TIP-network undergoes LLPS. This +TIP-droplet has then the potential to, e.g. co-condenses tubulin, and drive microtubule growth. Once the GTP-cap is hydrolyzed, the +TIP-droplet dissolves. This implies that the shape of the +TIP-droplets would follow the decaying GTP-tubulin profile at the growing microtubule tip. Whether the stoichiometry between EB3 and CLIP-170 has an impact on the LLPS process, as well as the distinct fluid properties and dynamics of an EB3/CLIP-170 droplet, merit investigation in future work.

A EB3/FL-CLIP droplet has distinct surface properties that differ from EB3- or FL-CLIP droplets in: surface tension, mobile fraction and tubulin interaction. These characteristics might be important for microtubule - EB3/FL-CLIP droplet interaction. Interestingly, we observed that EB3/FL-CLIP droplets on microtubules strongly enriched tubulin, while tubulin was less condensed in free floating EB3/FL-CLIP droplets. This might indicate that the microtubule surface can change the propensity of the EB3/FL-CLIP droplets to condense tubulin.

We hypothesize that LLPS of +TIPs serves an organizational purpose, allowing for the formation of a highly concentrated protein network at microtubule ends. Based on our observation that +TIPs concentrate soluble tubulin into droplets and on microtubules, we postulate that this process may contribute to regulation of microtubule dynamics in cells. Interestingly, depletion of CLIP-170 or EB3 from cells has been reported to only mildly affect microtubule dynamics in cells (Komorova et al., 2002), while depletion of EB3 and CLIP-170 reduced the growth speed by 20 % (Figure S7) Given that there is a level of redundancy in the functions of EBs in regulating microtubule dynamics in cells (Komorova et al., 2009; Yang et al., 2017), and the observation that diverse +TIPs partition into CLIP-170 condensates (Wu et al., 2021), we favor the hypothesis that +TIP-network condensation can be driven by multiple different proteins. In this scenario, individual depletions of CLIP-170 or EB3 would not strongly diminish microtubule growth, as other +TIPs could compensate for driving +TIP-network phase separation. Further studies comparing the phase-separation potencies of other +TIPs, as well as combinatorial depletions of LLPS-potent +TIPs in cells, will be of interest in addressing this question.

Is phase separation a common feature of +TIPs? Studies performed in parallel to our work show that this phenomenon is conserved across evolution: +TIPs in budding yeast, fission yeast, and higher eukaryotes have recently been demonstrated to undergo phase separation (Maan et al., 2021; Meier et al., 2021; Song et al., 2021, Jijumon et al., 2022). Intriguingly, in line with our results, the yeast studies confirmed that the CLIP-170 homolog played a key role in the phase separation process, whereas LLPS potency of EB homologs varied between organisms. The role of different mammalian EB family members in regulating LLPS will be an interesting direction for future studies. The ability of +TIP-networks to phase separate depends on intrinsically disordered regions (Maan et al., 2021; Song et al., 2021) and multivalent interaction modules (Meier et al., 2021), consistent with the observation that these features are highly evolutionarily conserved across +TIPs (Wu et al., 2021). Further studies will be necessary to investigate whether additional +TIPs contribute to the formation and regulation of +TIP-droplets.

Our work here and recent studies demonstrate that +TIP networks can behave like liquid condensates (Wu et al., 2021; Maan et al., 2021; Meier et al., 2021; Jijumon et al., 2022; Song et al., 2021). This work adds to the growing list of microtubule-related processes that are driven by LLPS and provides an exciting new paradigm for how cells can spatiotemporally control microtubule dynamics through local tubulin concentration (Jiang et al., 2015; Woodruff et al., 2017; Hernández-Vega et al., 2017; King and Petry, 2020; Jiang et al., 2021; Maan et al., 2021; Meier et al., 2021; Song et al., 2021). Interrogating the mechanical properties and composition of +TIP-droplets, as well as studying their regulation throughout the cell cycle, will be exciting avenues for future research.

## Acknowledgements

We thank Thomas Surrey (Centre for Genomic Regulation, Barcelona, Spain) for providing the FL-CLIP-170 expression vector for insect cells, and Michel Steinmetz (Paul Scherrer Institute, Villigen PSI, Switzerland) for providing vectors for EB3 purification. We would also like to thank Peter Bieling (Max Planck Institute of Molecular Physiology, Dortmund, Germany) for helpful discussions regarding expression and purification of FL-CLIP-170, and Maria Hondele (University of Basel, Switzerland) for helpful discussions for phase separation experiments. We would like to thank Dimitri Moreau and Stefania Vossio from the ACCESS Geneva high-content microscopy facility for help with microscopy and data analysis. We also thank Oscar Vadas and Rémy Visentin of the Protein Platform (Faculty of Medicine, University of Geneva, Switzerland) for support in FL-CLIP-170 expression. We would also like to thank Karsten Kruse and Marcos Gaitan-Gonzales and for critically reading the manuscript.

JM has been supported by the SNSF, 31003A_182473; RTW has been supported by the NCCR Chemical Biology program; CA has been supported by the DIP of the Canton of Geneva, SNSF (31003A_182473), and the NCCR Chemical Biology program.

## Author Contributions

JM and RTW performed and designed the experiments with the help of CA. MCV purified the proteins. JM, RTW, and CA analyzed data. RTW and CA wrote the manuscript.

## Supplementary Figures S1-S6

**Figure S1:**
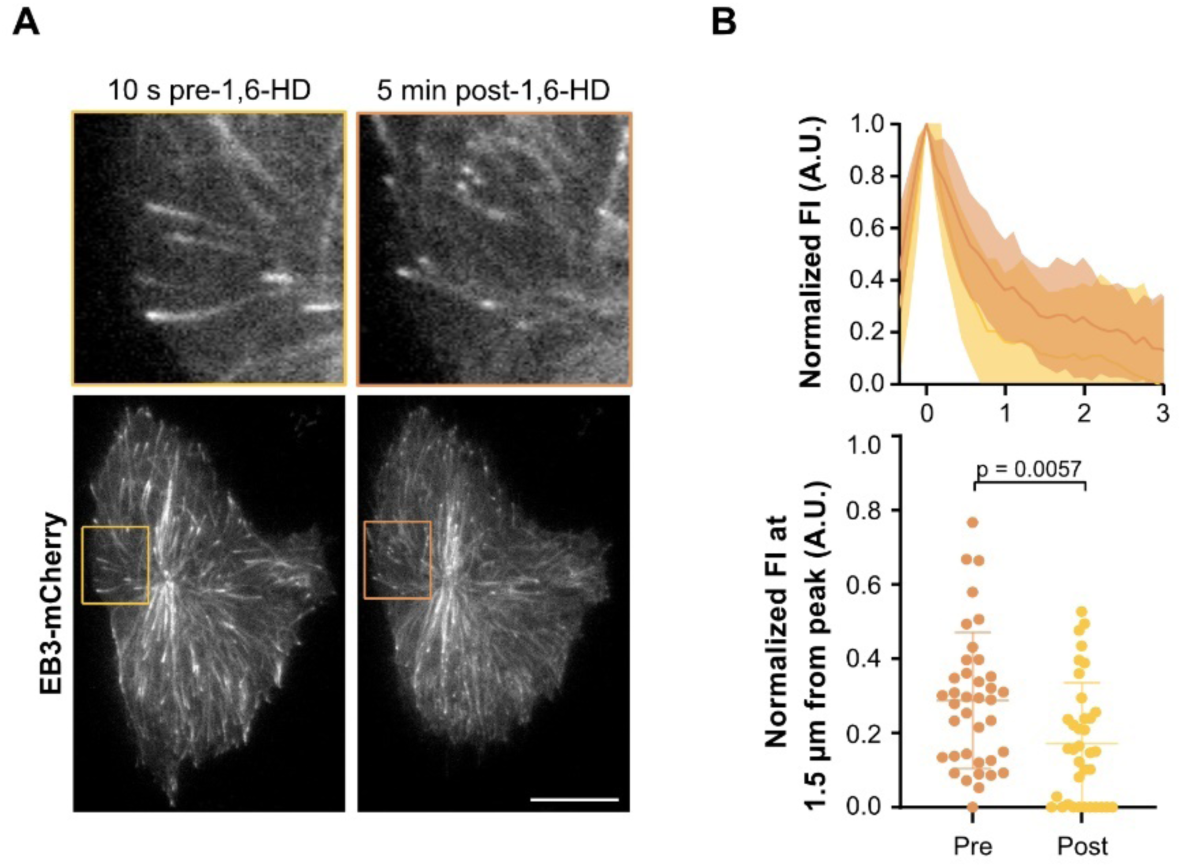
Decrease in EB3 network size upon treatment with 1,6-HD. **(A)** Representative images of CRISPR/Cas9 knock-in RPE-1-GFP-Tubulin cells expressing EB3-mCherry before and after 5-minute treatment with 5 % 1,6-hexanediol, with insets. Scale bar: 20 µm. **(B)** Top: mean fluorescence intensity profile of +TIP-networks before and after 1,6-hexanediol treatment. Bottom: normalized fluorescence intensity of +TIP-networks 1.5 µm away from the peak. Mean with SD of 36-38 +TIP-networks from 6 cells from 2 independent experiments. Statistics: paired t-test. Note that CLIP-170 patches were not dissolved by 1,6-hexanediol treatment.

**Figure S2:**
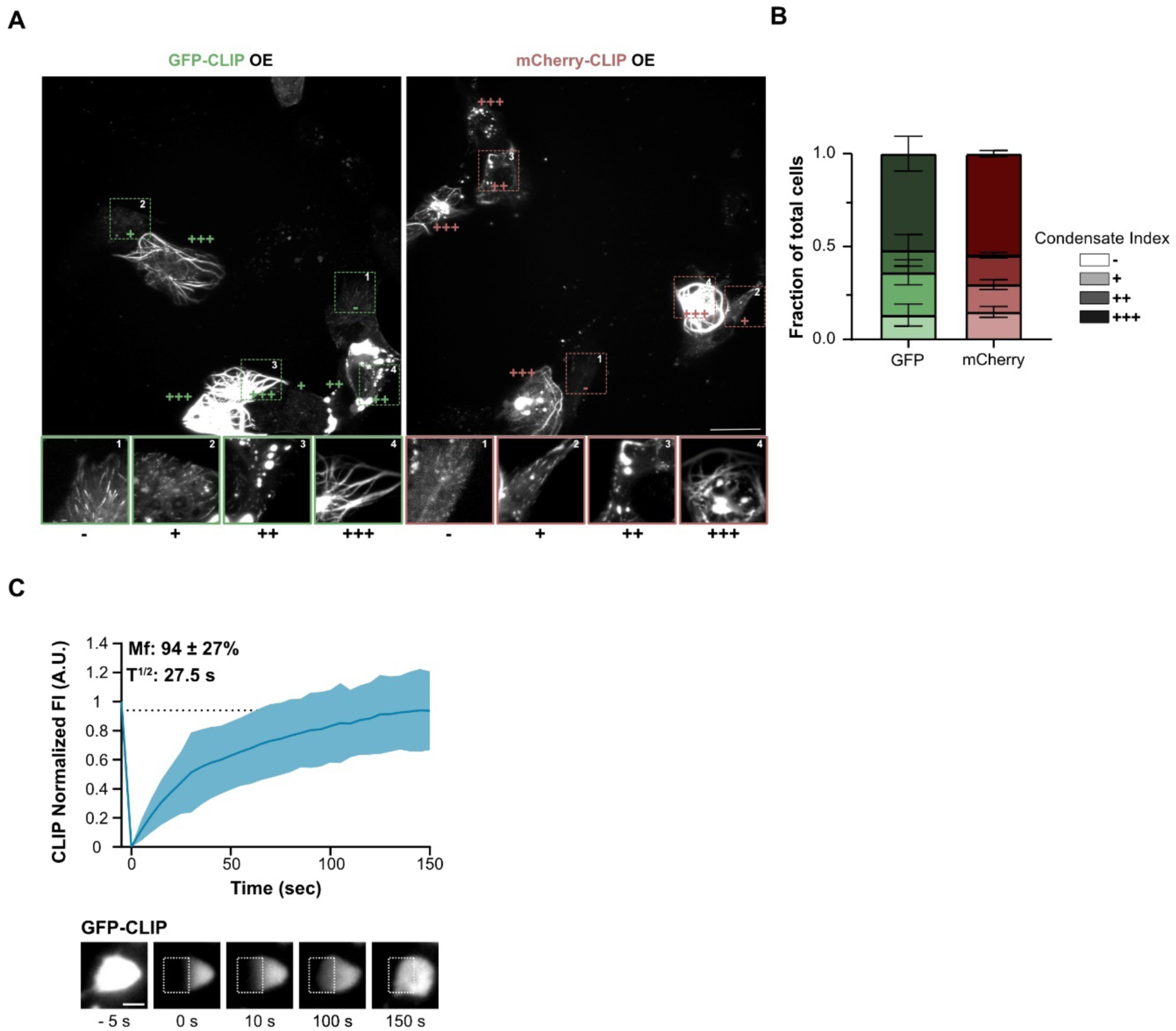
Fluorescent tag do not impact on LLPS of CLIP-170 in cells. **(A)** Representative images (top) with zoom-in (bottom) of RPE-1 cells expressing GFP-(left) or mCherry-CLIP (right) with indicated index for different condensation phenotypes. -: no observed cytoplasmic condensates; +: few small cytoplasmic condensates; ++: several large or many small condensates; +++: many large condensates and/or coating/bundling of microtubules. Note that zoom-in are contrast-adjusted. Scale bar: 30 µm. **(B)** Quantification of condensation index from experiments shown in A. Mean with SD from 2 independent experiments with 100 cells per experiment. (**C**) Representative TIRF images (bottom) and recovery curve (top) of GFP-CLIP-170 droplets in RPE-1 cells after photobleaching (dashed box). Mean with SD of 3 individual experiments with a total of 36 droplets from 24 cells. Scale bar: 2 µm.

**Figure S3:**
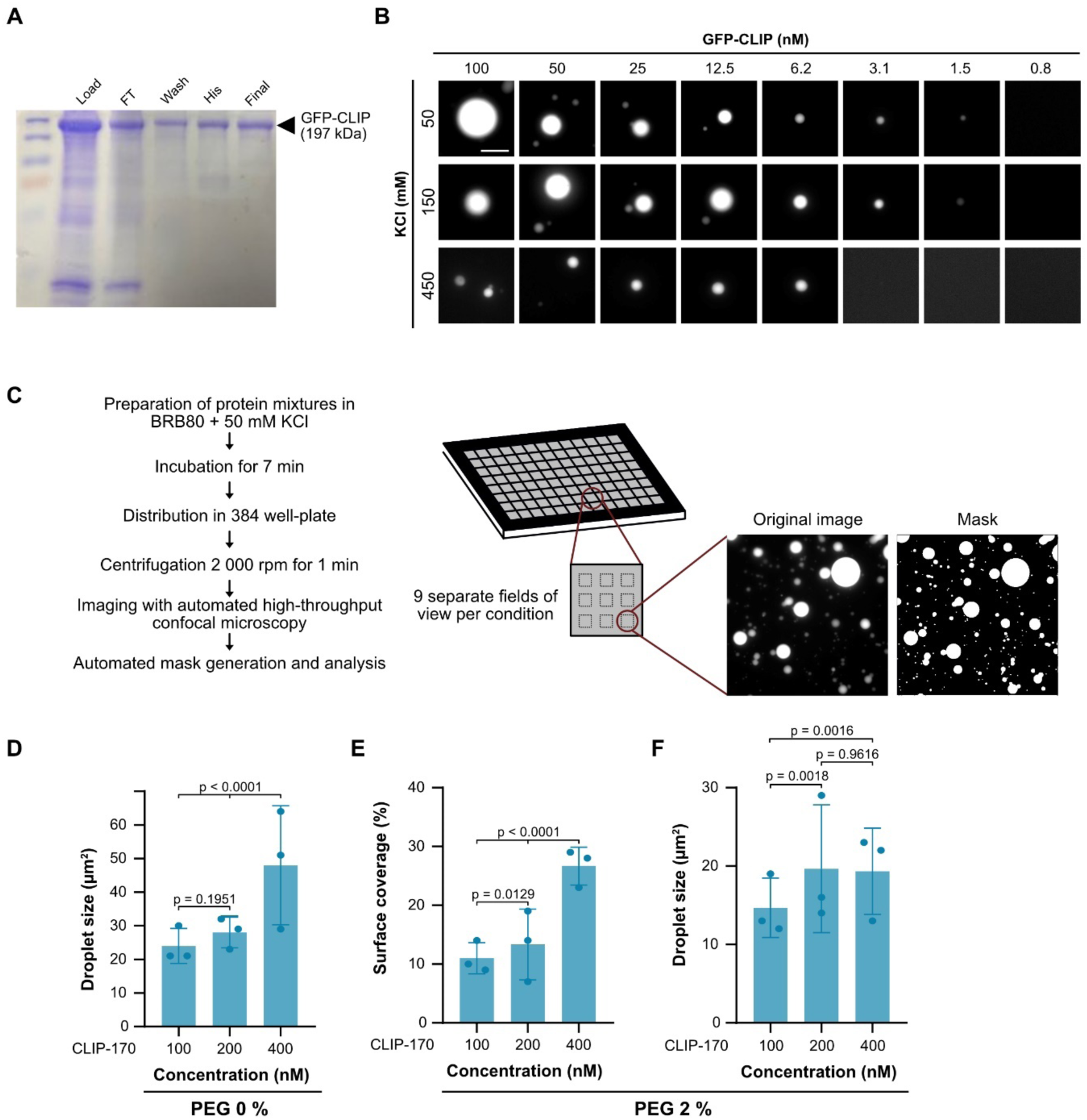
CLIP-170 droplet formation *in vitro* depends on molecular crowding, protein and salt concentrations. (**A**) SDS-PAGE analysis of GFP-FL-CLIP protein purification. Load, lysate loaded onto HisTRAP column; FT, flow through; wash, fractions collected during HisTRAP-column washing; final, final concentrated protein sample post-SEC and post-concentration. (**B**) Representative images of GFP-FL-CLIP phase diagram for 50, 150 and 450 mM KCl at indicated protein concentrations from 3 independent experiments. Scale bar: 10 µm. (**C**) Experimental outline for high throughput phase separation assays. For details, see materials and methods. (**D**) Droplet size (area) for GFP-FL-CLIP condensates at 100, 200, or 400 nM in the absence of PEG. (**E**) Coverslip surface coverage of GFP-FL-CLIP at the indicated concentrations in the presence of 2% PEG. (**F**) Droplet size (area) of GFP-FL-CLIP condensates at 100, 200 and 400 nM in the presence of 2 % PEG. Mean with SD from 3 independent experiments with a total of 27 fields of view. Statistics: two-tailed Student’s *t*-test.

**Figure S4:**
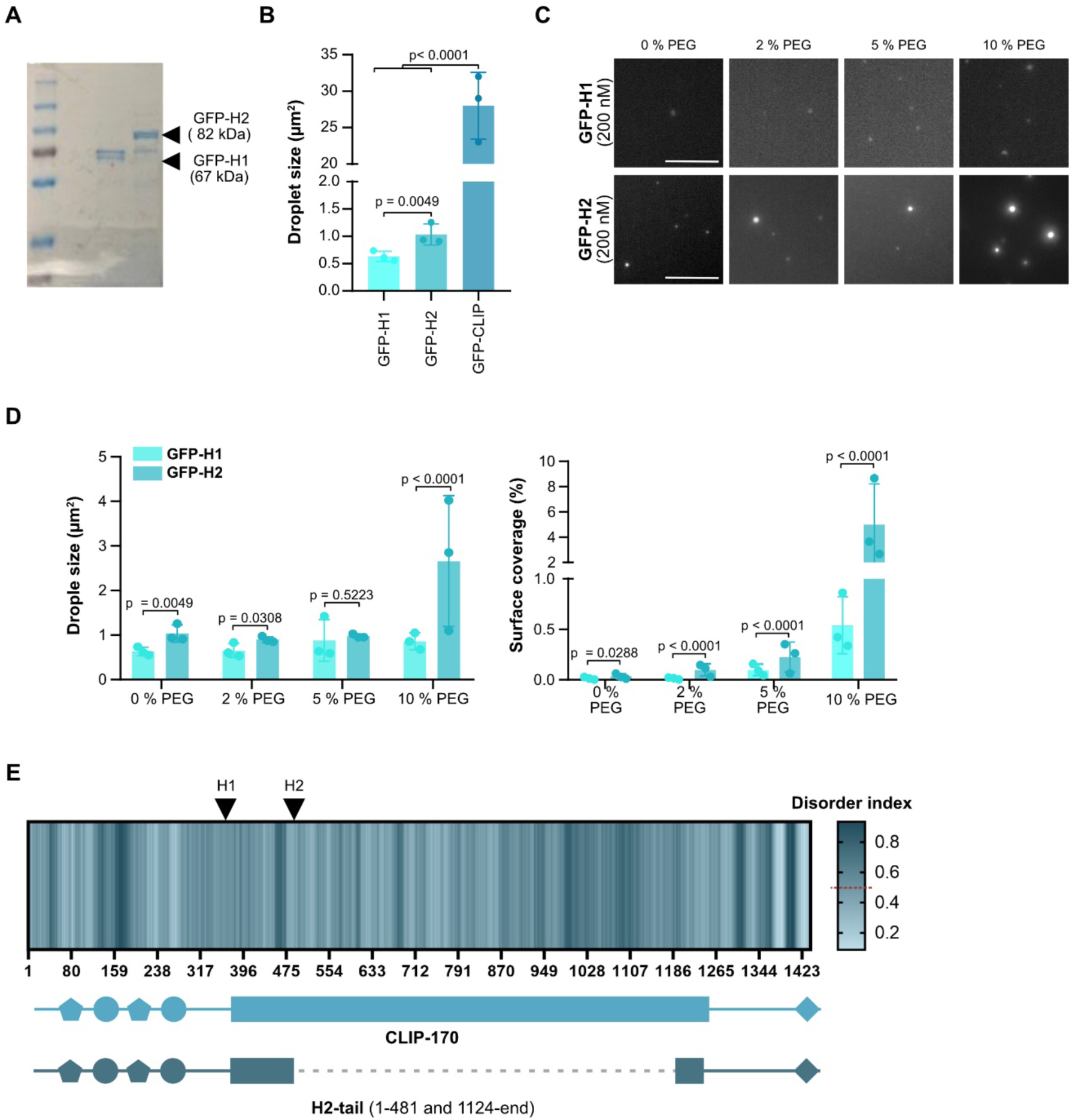
H2 only weakly undergoes LLPS even under strong molecular crowding conditions. (**A**) SDS-PAGE analysis of purified GFP-H1 and GFP-H2 protein. (**B**) Droplet size (area) of GFP-H2 (200 nM) and GFP-FL-CLIP (200 nM), and aggregate size of GFP-H1 (200 nM) in the absence of PEG. Mean with SD from 3 independent experiments with a total of 27 fields of view. Statistics: one-way ANOVA test. (**C**) Representative fluorescence confocal images of A in the presence of 0, 2, 5 and 10 % PEG. Scale bar: 20 µm. (**D**) Droplet size (left graph) and surface coverage (right graph) of denoted proteins at indicated PEG concentrations. Mean with SD of from 27 fields of view from 3 independent experiments. Statistics: two-tailed Student’s *t*-test. **(E)** Top: prediction of intrinsic disorder regions in CLIP-170, based on IUPred2A software. Values above the red dotted line (0.5) are considered as disordered (Mészáros et al., 2018). Dark blue corresponds to highly disordered regions. Bottom: CLIP-170 and mutant H2-tail secondary structures, length of H1 and H2 is indicated, black arrowhead.

**Figure S5:**
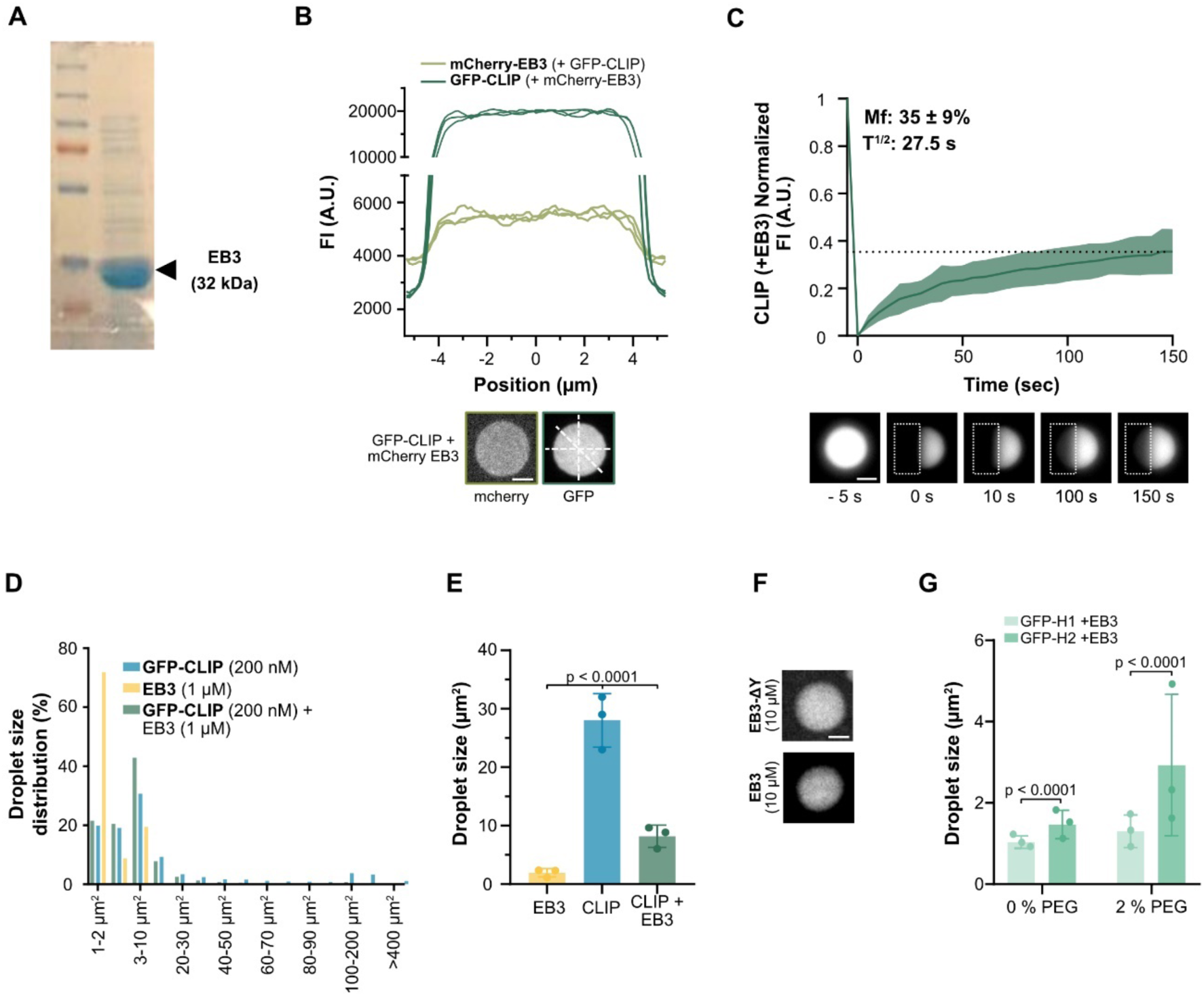
EB3/CLIP-170 phase separation is driven by the CLIP-170 C-terminal region *in vitro.* (**A**) SDS-PAGE analysis of purified EB3. (**B**) Line scan (top) with corresponding images (bottom, white lines indicate the line scans) of EB3/FL-CLIP droplets showing a homogenous protein distribution. GFP-FL-CLIP (2 µM) EB3 (8 µM unlabeled EB3 + 2 µM mCherry-EB3). (**C**) Representative images and recovery curve of purified EB3/GFP-FL-CLIP (10 µM / 2 µM) droplets after photobleaching (dashed box). Mean with SD of 3 independent experiments with a total of 35 condensates. Scale bar: 5 µm. (**D**) Size distribution of GFP-FL-CLIP (200 nM), EB3 (1 µM) and the EB3/FL-CLIP droplets in the absence of PEG. Mean size distribution from 3 independent experiments with a total of 27 fields of view. (**E**) Droplet size (area) of unlabeled EB3 (1 µM), GFP-FL-CLIP (200 nM) and EB3/FL-CLIP (1 µM + 200 nM) in absence of PEG. Mean with SD of 3 independent experiments with a total of 27 fields of view. Statistics: two-tailed Student’s *t*-test. (**F**) Representative images of EB3-Λ1Y condensates (8 µM unlabeled and 2 µM mCherry-EB3-Λ1Y) and EB3 (8 µM unlabeled and 2 µM mCherry-EB3). Scale bar: 4 µm. (**G**) Condensate size (area) of GFP-H1 (200 nM) or GFP-H2 (200 nM) in the presence of EB3 (1 µM) in the absence or presence of 2% PEG. Mean with SD of 3 independent experiments with a total of 27 fields of view. Statistics: two-tailed Student’s *t*-test.

**Figure S6:**
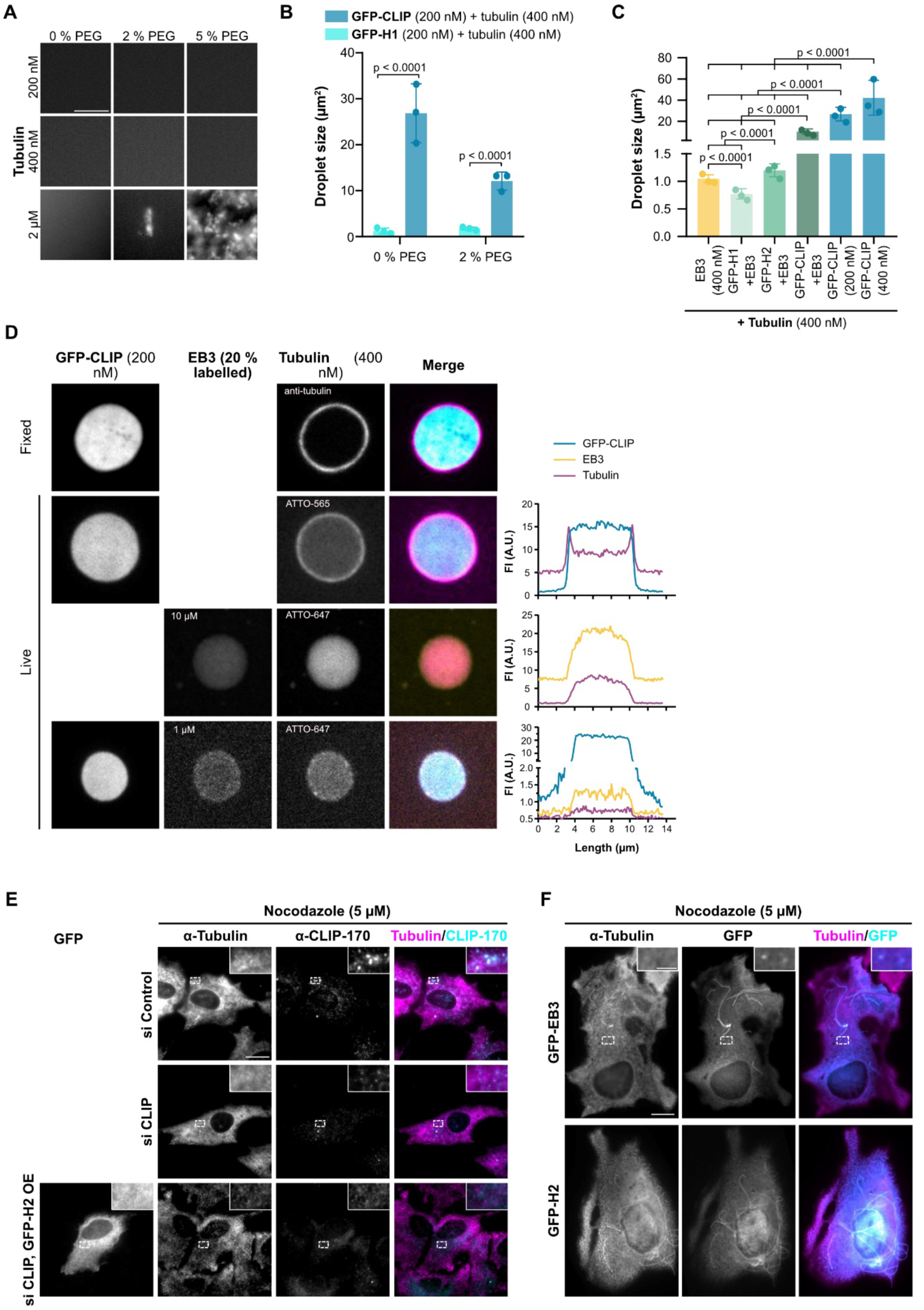
CLIP-170 and EB3 form a tubulin-condensing network. **(A)** Representative fluorescence confocal images of purified Atto-565-tubulin at indicated concentrations in presence of 0, 2 and 5 % PEG. Note that at 2 µM tubulin, PEG caused aggregation (but not condensate formation) of tubulin. Scale bar: 20 µm. (**B**) Quantified droplet size of Atto-565-tubulin (400 nM) in the presence of GFP-H1 (200 nM) or GFP-FL-CLIP (200 nM) in the absence or presence of 2% PEG. Mean with SD of from 27 fields of view from 3 independent experiments. Statistics: two-tailed Student’s *t*-test (right graph). (**C**) Quantified droplet size (area) of Atto-565-tubulin (400 nM) in the presence of purified EB3 (1 µM), GFP-H1 (200 nM) and EB3 (1 µM), GFP-H2 (200 nM) and EB3 (1 µM), GFP-FL-CLIP (200 nM) and EB3 (1 µM), GFP-FL-CLIP (200nM) alone, and GFP-FL-CLIP (400nM) alone. Mean with SD of three independent experiments. Statistic: two-tailed Student’s *t*-test. (**D**) Representative images of tubulin/FL-CLIP droplets (unlabeled tubulin) fixed and immunostained using tubulin specific antibodies (top). Representative images of GFP-FL-CLIP (200 nM) + tubulin-565 (400 nM) droplets, EB3 (8 µM) + mCherry-EB3 (2 µM) + tubulin-647 (4 µM) droplets and EB3 (900 nM) + mCherry-EB3 (100 nM) + GFP-FL-CLIP (200 nM) + tubulin-647 (400 nM) droplets with the corresponding line scans. Scale bar: 5 µm. (**E**) Representative images of fixed RPE-1 cells transfected with control siRNA (top panel), CLIP-170 siRNA (middle panel), or CLIP-170 siRNA rescued with GFP-H2 (bottom panel) treated with 5 µM nocodazole for 1 hour and stained for tubulin. Zoom-in of regions indicated by dashed box. Scale bar: 10 µm. (**F**) Representative images of fixed RPE-1 cells transfected with GFP-H2 (top panel) or GFP-EB3 (bottom panel) treated with 5 µM nocodazole for 1 hour and stained for tubulin. Zoom-in of regions indicated by dashed box Scale bar: 10 µm.

**Figure S7:**
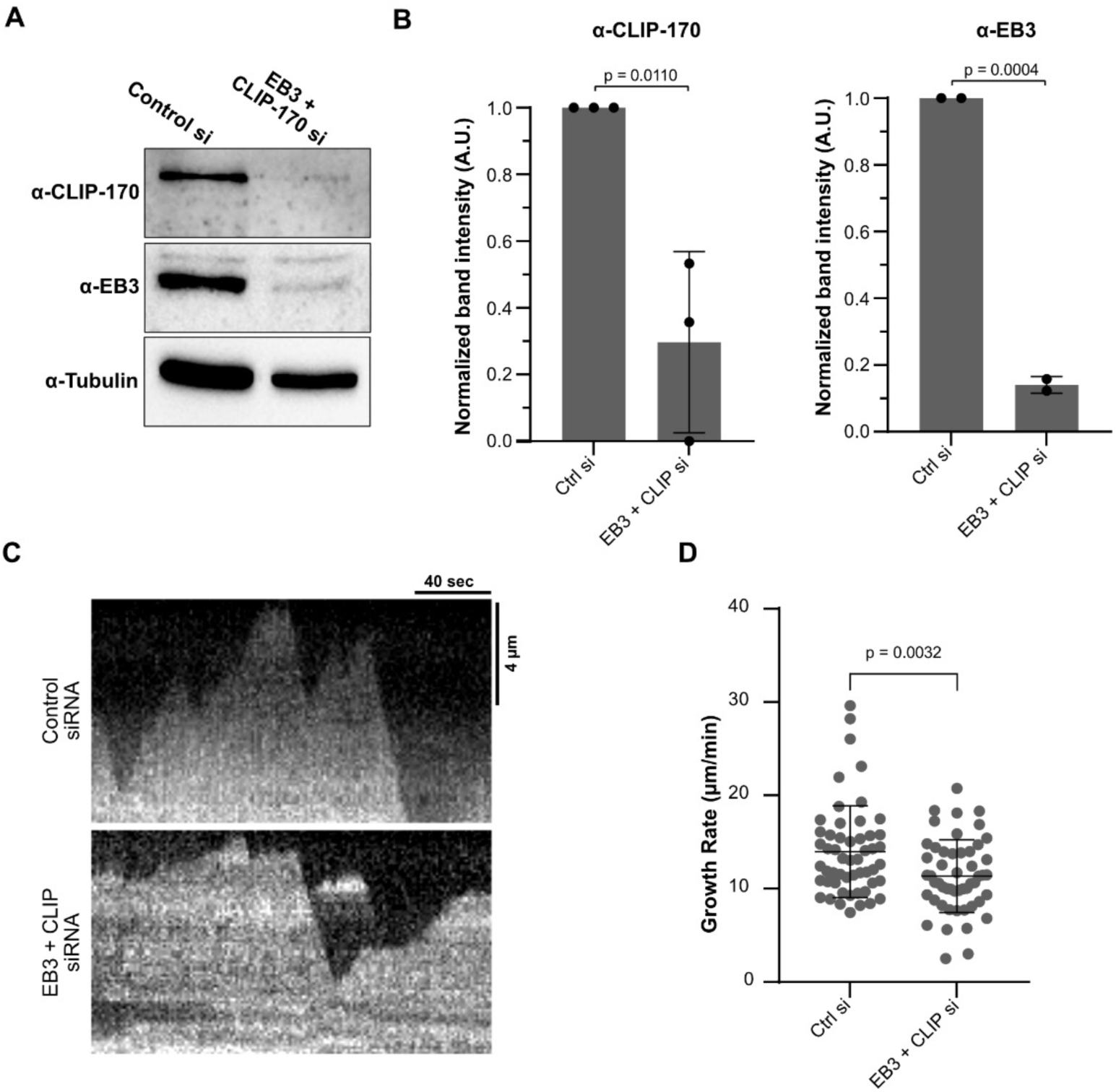
Depletion of EB3/CLIP-170 networks reduce microtubule growth rates in RPE-1 cells. (**A**) Representative Western Blot of CRISPR/Cas9 knock-in GFP-Tubulin RPE-1 cells transfected with control siRNA or siRNA to CLIP-170 and EB3 simultaneously for 72 hours. (**B**) Quantification of western blot of CLIP-170 depletion (left) and EB3 depletion (right) in cells treated with either control or EB3 + CLIP-170 siRNAs. Graphs show mean with SD from 3 (α-CLIP-170) or 2 (α-EB3) individual experiments. Statistics: paired t-test.(**C**) Representative microtubule kymographs from CRISPR/Cas9 knock-in RPE-1-GFP-Tubulin cells transfected with either control (top) or EB3 + CLIP-170 (bottom) siRNAs for 72 hours. (**D**) Mean microtubule growth rate with SD from: Control – 53 microtubules from 29 cells; EB3 + CLIP siRNA – 51 microtubules from 19 cells (4 independent experiments per condition). In graph, each dot represents a single microtubule. Statistics: paired t-test.

**Figure S8:**
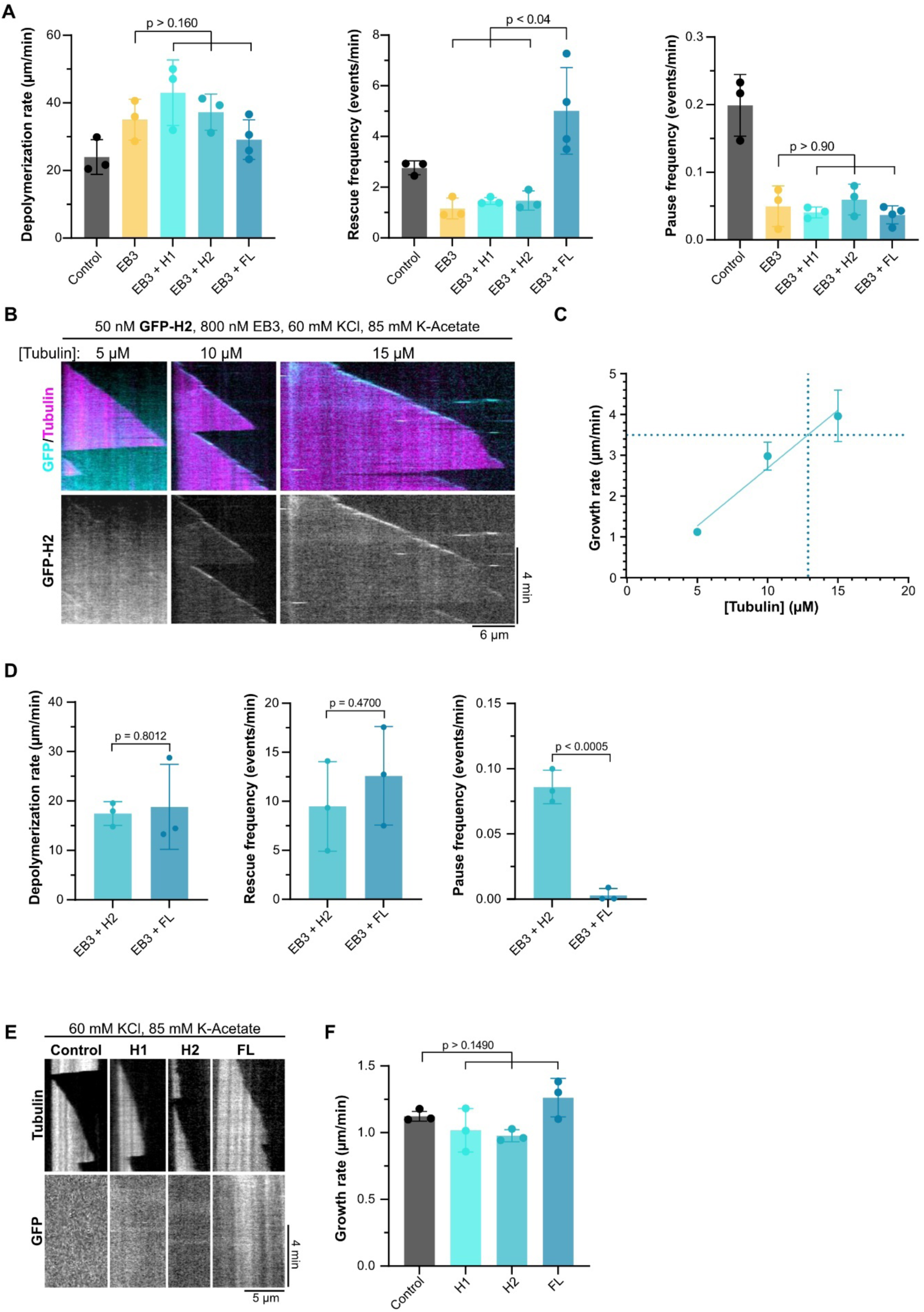
Phase separation potent +TIP-networks impart stronger effects on microtubule dynamics. (**A**) Microtubule dynamic parameters (depolymerization rate, left; rescue frequency, middle; and pause frequency, right) for the denoted conditions in 60 mM KCl and 85 mM K-acetate (corresponding assay to Figure 6B, see here for number of analyzed microtubules). Mean with SD of three individual experiments. Statistics: one-way ANOVA Fisher’s LSD test (**B**) Representative microtubule kymographs in the presence of GFP-H2 (50 nM) and EB3 (800 nM) grown at the denoted tubulin and salt concentrations. (**C**) Microtubule growth rate from experiments in B. Mean with SD from three individual experiments. Solid cyan line shows linear regression curve fit. Dashed blue line indicates concentration of tubulin at which microtubule in presence of EB3/H2 networks grow at 3.8 µm/min (the speed achieved by EB3/FL-CLIP droplets at 5 µM tubulin; Figure 6A). (**D**) Microtubule dynamic parameters (depolymerization rate, left; rescue frequency, middle; and pause frequency, right) for the denoted conditions in 60 mM KCl (corresponding assay to Figure 6H, see here for number of analyzed microtubules). Mean with SD from minimum three independent experimental replicates. Statistics: one-way ANOVA Fisher’s LSD test (**E**) Representative microtubule kymographs of control (only tubulin), 50 nM GFP-H1, 50 nM GFP-H2 or 50 nM FL-CLIP. (**F**) Microtubule growth rate from experiments in Figure E. Mean with SD from three independent experiments. Total number of MTs analyzed per condition: Control – 48; H1 – 42; H2 – 41; FL – 26. Statistics: one-way ANOVA Fisher’s LSD test.

**Figure S9:**
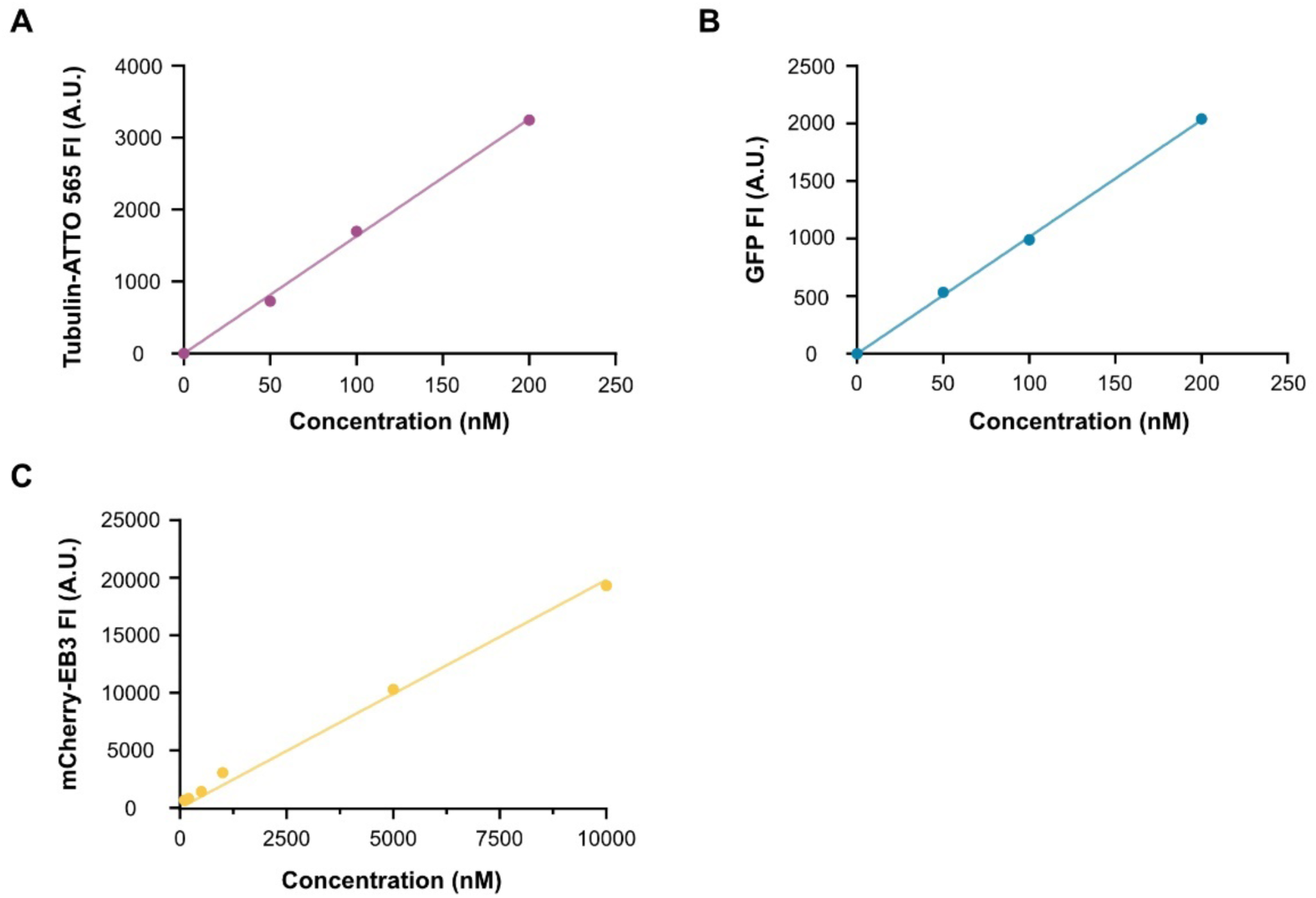
Calibration curve for Tubulin-ATTO 565, mCherry-EB3 and GFP. Calibration curve between fluorescent intensity and concentration of: (**A**) Tubulin-ATTO 565 at 50, 100 and 200 nM. (**B**) GFP at 50, 100 and 200 nM and (**C**) mCherry-EB3 at100, 200, 500, 5 000 and 10 000 nM.

**Movie 1:** Representative foci formation resembling fission in GFP-CLIP overexpressing RPE-1 cells. Scale bar: 2 µm.

**Movie 2:** Representative CRISPR/Cas9 knock-in GFP-tubulin RPE-1 expressing mCherry-EB3 and treated with 5% 1,6-hexanediol (added at 00:30).

**Movie 3:** Representative CRISPR/Cas9 knock-in GFP-tubulin RPE-1 expressing mCherry-CLIP. Inset highlights fusion of mCherry-CLIP droplets.

**Movie 4:** Representative CRISPR/Cas9 knock-in GFP-tubulin RPE-1 expressing mCherry-CLIP. Inset highlights fission of mCherry-CLIP droplets.

**Movie 5:** Fluorescence recovery after photobleaching of a mCherry-CLIP droplet in a representative CRISPR/Cas9 knock-in GFP-tubulin RPE-1 .

**Movie 6:** Fusion of two GFP-CLIP droplets (1µM) *in vitro*.

**Movie 7:** Fluorescence recovery after photobleaching of a purified GFP-CLIP droplet (2 µM) *in vitro*.

**Movie 8:** Representative CRISPR/Cas9 knock-in GFP-tubulin RPE-1 (magenta) expressing mCherry-H1 (cyan).

**Movie 9:** Representative CRISPR/Cas9 knock-in GFP-tubulin RPE-1 (magenta) expressing mCherry-H2 (cyan).

**Movie 10:** Fluorescence recovery after photobleaching of a purified EB3 (10 µM) labeled with GFP-EB3 (100 nM) droplet *in vitro*.

**Movie 11:** Fluorescence recovery after photobleaching of a purified EB3 (10 µM) labeled with GFP-EB3 (100 nM) mixed with GFP-CLIP (2 µM) droplet *in vitro*.

**Movie 12:** 3D representation of EB3 (8 µM) labelled with mCherry-EB3 (2 µM) droplets *in vitro*. Note that floating droplets moved during image acquisition.

**Movie 13:** 3D representation of GFP-CLIP (2 µM) droplets *in vitro*.

**Movie 14:** 3D representation of GFP-CLIP (2 µM) mixed with EB3 (8 µM) labelled with mCherry-EB3 (2 µM) droplets *in vitro*. Note that floating droplets moved during image acquisition.

**Movie 15:** Representative *in vitro* microtubule dynamics assay of Atto-565 tubulin (magenta) polymerized in the presence of 50 nM GFP-FL-CLIP (cyan) and 800 nM EB3 in high salt buffer.

**Movie 16:** Representative *in vitro* microtubule dynamics assay of Atto-565 tubulin (magenta) polymerized in the presence of 50 nM GFP-FL-CLIP (cyan) and 800 nM EB3 in low salt buffer.

## Methods

### Cell culture and treatments

Parental RPE1 and CRISPR/Cas9 knock-in RPE1-GFP-tubulin cells were cultured in high glucose Dulbecco’s Modified Eagle’s Medium F12 (DMEM, ThermoFisher, 113057) supplemented with 10 % Fetal Bovine Serum (FBS, ThermoFisher, 10270106) and 1 % penicillin-streptomycin (Gibco, 15140122) at 37°C with 5 % CO_2_. The cell lines were monthly checked for mycoplasma contamination. CRISPR/Cas9 knock-in GFP-tubulin RPE-1 cells were generated using the same guide RNA and protocol as in Andreu-Carbó et al., 2022.

For transient expression studies of exogenous FL-CLIP, H1, H2 and EB3, cells were transfected using the jetOPTIMUS transfection reagent (Polyplus) with 0.5 µg DNA according to the manufacturer’s instructions. Transfection media was replaced with fresh culture media 8 hours post-transfection, and cells were imaged 15-24h after transfection.

For experiments in which endogenous CLIP-170 was depleted, parental RPE1 cells were transfected with 10 nM siRNA targeting CLIP-170 (Santa Cruz Biotechnology) using lipofectamine RNAiMAX reagent (Invitrogen) according to the manufacturer’s instructions. Media was replaced the following morning, and cells were cultured for 72 hours post-transfection before fixation, or transfected with GFP-H2 24 hours pre-fixation for “knockdown-rescue” experiments with H2. For control experiments, cells were transfected with Allstars Negative Control siRNA (QIAGEN). For CLIP-170 + EB3 depletion experiments, cells were transfected with dual siRNA mixes targeting EB3 (Thermo Fisher, s22683) and CLIP-170 (Santa Cruz, 43281). Transfection media was replaced with fresh culture media 8 hours post-transfection, and cells were imaged 72 hours post-transfection.

To depolymerize the microtubule network, cells were treated with 5 µM nocodazole (Sigma, M1404; diluted in culture medium) for 1 hour prior to fixation. For experiments using 1,6-hexanediol (Sigma), cells were treated with 5% 1,6-hexanediol (diluted in culture medium) for 10 minutes.

### Cloning

The in-cell expression vector for mCherry-FL-CLIP170 was generated by excising GFP from a GFP-FL-CLIP170 vector (a kind gift from Thomas Surrey) using AgeI/BsrG1 restriction sites and replacing it with mCherry containing AgeI/BsrG1 overhangs generated by PCR. From this vector, we generated mCherry-tagged H1- and H2-CLIP170 by PCR and reinsertion using the XhoI/KpnI restriction sites (N-terminal XhoI primer: 5’-CCGCTCGAGCTCAAGCTTCGATGAGTATGCTGAAACCCAGCGGGCTGAA-3’, C-terminal KpnI H1 primer: 5’-CGGGGTACCGTCGACTCAAGTGGTGCCCGAGATCTTGCGGGC-3’, and C-terminal KpnI H2 primer: 5’-CGGGGTACCGTCGACTCATTTGTCAGCTTTGGTCTTTTCAAAGAGCAGGCTCTGTTC-3’). Protein purification vectors for GFP-H1- and H2-CLIP were generated by PCR from the GFP-FL-CLIP-170 vector using a primer with an overhang for the Ndel restriction site as well as an N-terminal TEV protease site (5’- GCGGCAGCCATATGGAAAACCTGTATTTCCAGGGAAGTGCCACCATGGTGAGCAAG GGCGAGGAGCTGTTCA-3’), and C-terminal primers specific to each CLIP truncation with overhangs corresponding to the ScaI restriction site (H1: 5’-CCTTATCAAGTACTAGTGGTGCCCGAGATCTTGCGGGCGTAGCGGGAAG-3’) (H2: 5’-CCTTATCAAGTACTTCATTTGTCAGCTTTGGTCTTTTCAAAGAGCAGGCTCTGTTCAAGC-3’), then cloned into an empty pET28a-6His vector (a kind gift from Natacha Olieric, Paul Scherrer Institute) using NdeI/ScaI restriction sites.

For the GFP-H2-tail construct Gilson assembly was used from the GFP-FL-CLIP construct using the primer 5’- CAAGCTTCGAATTCTATGCTGAAACCCAGCGGGCTGAAGG with BstBl (Start of H2), CAGCTCTGCGTCCTTCTCTTTGTCAGCTTTGGTCTTTTCAAAGAGCAG-3’ (end of H2), 5’- GCTGACAAAGAGAAGGACGCAGAGCTGGAGAAGCTGAGGAATGAG (start of tail) and CCGCGG TACCGTCGACTCAGAAGGTCTCATCGTCGTTGCAGTTGG (end of tail) using the PKPN enzyme.

A protein purification vector for mCherry-6His-EB3 was a kind gift from Natacha Olieric (Paul Scherrer Institute). From this vector, mCherry was excised using AgeI/BsrGI restriction sites to produce an untagged 6His-EB3 vector for purification. Mutagenesis of the mCherry-6His-EB3 was used to create the EB3-1′Y281, the primer used were 5’-CAAGAAGACCAGGACGAGTAACAGTAAAGGTGGATAC and GTATCCACCTTTACTGTTACTCGTCCTGGTCTT CTTG-3’.

For in-cell expression of EB3, EB3-mCherry vectors were obtained from Addgene (Addgene plasmid 55037).

### Imaging

#### Microscope

For in-cell studies and *in vitro* microtubule dynamics experiments, imaging was performed on an Axio Observer Inverted TIRF microscope (Zeiss, 3i) equipped with a Prime 95B BSI (Photometrics) using a 100X objective (Zeiss, Plan-Apochromat 100X/1.46 oil DIC (UV) VIS-IR). SlideBook 6 X 64 software (version 6.0.22) was employed to record time-lapse imaging. For *in vitro* microtubule dynamics and cell imaging, microscope stage conditions were controlled with the Chamlide Live Cell Instrument incubator (37°C for *in vitro* experiments, supplemented with 5 % CO_2_ for live cell experiments).

For stack acquisition and intensity measurement a 3i Marianas spinning disk confocal setup based on a Zeiss Z1 stand, a 100× PLAN APO NA 1.45 TIRF objective and a Yokogawa X1 spinning disk head followed by a 1.2× magnification lens and an Evolve EMCCD camera (Photometrics). Fast z-stack acquisition (0.5-µm steps) was obtained using a piezo stage (Mad City Labs). Single-emitter emission filters were always used to avoid bleed-through and each channel was acquired sequentially.

#### Microtubule dynamics

For *in vitro and in cell* microtubule dynamics measurements images were taken every second for 3 minutes. Microtubules were tracked individually using the Freehand-Line tool in ImageJ (15-pixel width) and kymographs were built using the KymographBuilder plugin. Microtubule growth speeds were then calculated by manually tracing the slopes of kymographs using the Straight-Line tool in ImageJ and extracting dynamic parameters the slopes using a custom-written code.

#### Fluorescence recovery after photobleaching

Fluorescence recovery after photobleaching (FRAP) experiments in cells were performed in square regions (4x4 µm) with a 656 nm laser at 20% intensity. The normalized fluorescence intensity was calculated using the formula 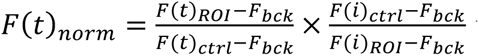 where F(t)_ROI_ and F(t)_ctrl_ are respectively the ROI and the control fluorescence intensity before the FRAP, F_bck_ the background fluorescence and F(i)_ROI_ and F(i)_ctrl_ are respectively the ROI of the unbleached part of the condensate at one timepoint (i) (Day et al.,2012).

#### +TIP-network analysis in 1,6-hexanediol-treated cells

For experiments in cells treated with 1,6-hexanediol, images were taken in a single z-plane every 10 seconds for 10 minutes. 1,6-Hexanediol (5 %) was added after one minute, and cells were only analyzed if they did not undergo any large-scale changes in morphology, as 1,6-hexanediol treatment has been noted to affect mammalian cell shape (Wheeler et al., 2016). For “pre-treatment” time points, all in-focus +TIP-networks on the cell periphery were analyzed in the time frame 10 seconds prior to hexanediol addition using the Segmented-Line tool in ImageJ to obtain fluorescence intensity. For “post-treatment” time points, the same strategy was applied to the time frame 5 minutes after hexanediol addition.

### Immunofluorescence

15-24 hours post-transfection (or post-seeding for non-transfected cells), cells were fixed with 100% methanol for 5 min at -20°C and then for 15 min with 3% paraformaldehyde at room temperature. Cells were then permeabilized for 10 minutes with 0.15% by volume Triton-X 100 (Sigma) in PBS followed by 10 minutes with 0.1% Tween-20 (AppliChem) in PBS, washed thoroughly in a solution of 0.05% Tween-20 in PBS (hereafter referred to as PBS-T), and subsequently blocked with 2% bovine serum albumin (in PBS) for 1h. Post-blocking, cells were incubated overnight with antibodies targeting tubulin (Sigma T6199, DM1α, 1:1000, mouse), EB1 (Millipore AB 6057, rabbit, 1:1000), EB3 (Santa Cruz Biotechnology sc-101475, KT36, rat, 1:200), or CLIP-170 (Santa Cruz Biotechnology sc-28325, F3, mouse, 1:500). Primary antibodies were diluted to the appropriate concentration in 2% bovine serum albumin in PBS. The following day, cells were subjected to three five-minute washes at room temperature in PBS-T, then subsequently incubated in secondary antibodies (Invitrogen, species-specific IgG conjugated to Alexa-647, 568, or 488 fluorophores) at room temperature for one hour. Cells were subjected to three additional PBS-T washes, and coverslips were mounted onto glass microscopy slides (Glass technology) using ProLong^TM^ Diamont Antifade Mountant. Coverslips were sealed with nail polish and stored at 4°C until imaging.

### Tubulin purification from bovine brain and labelling

Tubulin was purified from fresh bovine brain by two subsequent polymerization/depolymerization cycles as described previously (Andreu-Carbó et al., 2022). Tubulin labelling with biotin or ATTO-488, -565, -647 fluorophores was performed as described (Andreu-Carbó et al., 2022), and final labelling ratios to polymerize microtubules were 11% for ATTO-488 and 13% for ATTO-565 tubulin.

### Protein purification

For purification of EB3, mCherry-EB3, EB3-ΔY and mCherry-EB3-ΔY, *E. coli* BL21 (DE3) cells were transformed with 6-His-tagged EB3-encoding plasmids and induced for expression overnight with 1 mM IPTG at 20°C under rotation at 200 rpm. All following steps were performed at 4°C. The morning after induction, cells were lysed by in lysis buffer (20 mM Tris pH 7.5, 300 mM NaCl) supplemented with 1% Triton-X 100 and protease inhibitors cocktail tablets (Roche) and sonicated. Cell debris were then cleared by ultracentrifugation. The cleared lysate was subsequently loaded onto a pre-equilibrated HisTrap column (GE Healthcare 1mL HisTrap column) using an ÄKTA Pure Protein Purification System (GE Healthcare). After washing the column in lysis buffer, elution buffer (20 mM Tris pH 7.5, 300 mM NaCl, 1 M imidazole) was applied to the column in a 1% gradient. Eluted protein fractions were pooled and concentrated using Amicon 30K Centrifugal filters (Millipore). The concentrated, cleared protein was subjected to size-exclusion chromatography using a HiLoad 16/600 Superdex column (GE Healthcare) in lysis buffer. Protein-containing fractions were harvested, pooled, and concentrated. Protein was supplemented with 20% glycerol, aliquoted, snap-frozen and stored at -80°C.

GFP-H1, GFP-H2 and GFP-H2-tail were purified using the same scheme as EB3, with the following differences: (1) the lysis buffer was 50 mM potassium phosphate pH 7.5, 500 mM NaCl, 1 mM MgCl_2_, and 1 mM β-mercaptoethanol. (2) Between applying protein to the HisTrap column and elution, the column was washed with lysis buffer supplemented with 8 mM Imidazole. (3) H1- and H2-CLIP were eluted from the HisTrap column using lysis buffer + 300 mM Imidazole, and protein-containing fractions were subjected to tobacco etch virus (TEV) protease treatment overnight to remove His tags prior to size-exclusion chromatography.

FL-CLIP170-GFP was purified from insect cells as described previously (Telley et al., 2011). A plasmid encoding FL-CLIP170-GFP in pFasBacHTa (a kind gift from Thomas Surrey) was used to generate Baculovirus, which was subsequently used to infect Sf9 cells. Cells were harvested and lysed with lysis buffer (30 mM HEPES pH 7.4, 400 mM KCl, 20 mM Arginine, 20 mM potassium-glutamate, 0.01% Birj35, 2 mM MgCl_2_, 10 mM β-mercaptoethanol) supplemented with 20 mM imidazole and protease inhibitor tablets (Roche) using dounce homogenization. Cell debris were cleared by ultracentrifugation, and cleared lysate was loaded onto a Histrap column (GE Healthcare). The column was washed with lysis buffer supplemented with 50 mM imidazole, then eluted with lysis buffer supplemented with 300 mM imidazole. Protein-containing fractions were pooled and further cleared by a second centrifugation, then subjected to size-exclusion chromatography using a HiLoad 16/600 Superdex column (GE Healthcare) in lysis buffer (lacking Birj35). Only proteins coming off with a clean, single peak profile from the Superdex column were used. Protein-containing fractions were pooled and concentrated and used immediately (maximal 5 hrs after purification), as FL-CLIP was prone to degradation and loss of activity after freezing as previously noted (Telley et al., 2011). For all purified proteins, protein concentration was measured by Bradford assay.

### Phase separation assay

For in vitro phase separation assays, proteins were diluted to the appropriate concentration in BRB80 supplemented with 50 mM potassium chloride and PEG 4000 (0, 2, 5 or 10 % by weight) in Eppendorf tubes. After thorough mixing, reactions were transferred to 384-well plates (Falcon) and incubated for 7 min. The plate was then centrifuged at 2000 rpm for 1 min to sediment proteins in the dense phase on the well bottoms. Images were acquired using a confocal automated microscope (Molecular Device) with a 60X dry objective. For each reaction, 9 times 2048 x 2048 px fields of view were acquired with one focal plane (Figure S1A). Automated analysis was performed using MetaXpress Custom Module editor software. From fluorescence intensity, masks were generated to differentiate condensates from the background and the area sum was calculated together with the condensate versus background fluorescence intensity.

### Droplet-Pelleting assay

For the droplet-pelleting assays to determine amount of protein in the dense phase (Figure 5A) proteins were diluted to the appropriate concentrations (1 µM CLIP, 10 µM EB3, or 1 µM CLIP + 10 µM EB3) in BRB supplemented with 125 mM potassium chloride and incubated at room temperature for 30 minutes. Following this incubation, reaction mixtures were centrifuged at 16,900 g at room temperature for 15 minutes using a Fresco 21 Haraeus tabletop centrifuge (Thermo Scientific). The supernatant was collected, and pellets were resuspended in an equal volume of resuspension buffer (BRB80 + 125 mM KCl). Pellets and supernatants from each reaction were run on SDS-PAGE gels, which were subsequently stained with QuickBlue Protein Stain Coomassie dye (Lubio Science) for at least 2 hours before destaining in water. The fraction of protein in dense vs dilute phase was calculated by dividing the integrated Coomassie band intensity from the pellet or supernatant, respectively, by the sum of the integrated band intensity of the pellet and supernatant. Fold-change in pellet fraction was taken by dividing the protein fraction in the pellet for experiments with CLIP + EB3 by the protein fraction in the pellet for each protein alone.

### Fixation of FL-CLIP droplets

For experiments in which FL-CLIP droplets were fixed to observe whether unlabeled tubulin partitions into droplets (Figure S6B), GFP-CLIP (200 nM) was incubated with unlabeled purified bovine brain tubulin (400 nM) on coverslips for 15 minutes. Reactions were fixed directly on coverslips with 3% PFA for 15 minutes, washed 3 times in PBS, blocked and stained with antibodies to tubulin as described in the above immunofluorescence procedure.

### Calibration curve

To measure the concentration of proteins in the dilute and in the droplet phase, calibration curves were established for GFP, tubulin-565 and mCherry-EB3. To avoid mCherry-EB3 phase separation a buffer with 1M NaCl was used. To establish the curve the following concentrations were used: GFP (50, 100 and 200 nM); mCherry-EB3 (100, 200, 500, 5 000 and 10 000 nM); and tubulin (50, 100 and 200 nM).

### Coverslip treatment and Flow chamber preparation

For *in vitro* microtubule dynamics studies, slides and coverslips were cleaned by two successive 30-minute sonication cycles in 1 M NaOH followed by 96% ethanol with thorough rinsing in bi-distilled water between each step. After drying, slides and coverslips were plasma treated (Electronic Diener, Plasma surface technology) and subsequently incubated for 48 hours with tri-ethoxy-silane-PEG (Creative PEGWorks) or a 1:5 mix of tri-ethoxy-silane-PEG-biotin: tri-ethoxy-silane-PEG (final concentration 1 mg/ml) in 96 % ethanol and 0.02 % HCl, with gentle agitation at room temperature. Slides and coverslips were then washed in ethanol (96 %) followed by thorough washing in bi-distilled water, then dried with an air gun and stored at 4°C. Flow chambers were prepared by affixing a silane-PEG-biotin coverslip to a silane-PEG slide using double-sided tape.

### Microtubule dynamics assays in vitro

Microtubule seeds were prepared at a final concentration of 10 µM tubulin (20 % ATTO-647-labelled tubulin and 80 % biotinylated tubulin) in BRB80 supplemented with 0.5 mM GMPCPP (Jena Bioscience) for 45 minutes at 37°C. Seeds were incubated with 1 µM Paclitaxel (Sigma) for 45 minutes at 37°C, centrifuged (50,000 rpm at 37°C for 15 min), resuspended in BRB80 supplemented with 1 µM Paclitaxel and 0.5 mM GMPCPP, aliquoted and subsequently stored in liquid nitrogen.

Flow chambers were prepared by injecting subsequently 50 µg/mL neutravidin (ThermoFisher), BRB80, and microtubule seeds, then subsequently washing out unattached seeds with BRB80. Reaction buffer containing Atto-565 labelled-tubulin (1:5 ratio labelled to unlabeled; 5 µM for all assays except for Figure S6B, C) in BRB80 supplemented with an anti-bleaching buffer [10 mM DTT, 0.3 mg/mL glucose, 0.1 mg/mL glucose oxidase, 0.02 mg/mL catalase, 0.125 % methyl cellulose (1500 cP, Sigma), 1 mM GTP] was subsequently injected, and chambers were sealed with silicon grease and immediately imaged. For “low salt” assays (Figure 6E-F), the reaction buffer was supplemented with 60 mM potassium chloride. For “high salt” assays (Figure 6A-B), the reaction buffer was supplemented with 60 mM potassium chloride and 85 mM potassium acetate to facilitate tip tracking as described previously (Telley et al., 2011).

For assays involving recombinant EB3 and H1- or H2-CLIP-170, purified proteins were flash-thawed and spun at 50,000 rpm in a TLA-100 centrifuge at 4°C for 15 minutes to remove any large aggregates. Proteins were diluted into BRB80 immediately prior to their usage, and further diluted to the appropriate concentration in reaction buffer. Assays involving FL-CLIP-170 were carried out as noted above, but within the first 5 hours post-purification as FL-CLIP-170 activity is poorly preserved after freezing (Telley et al., 2011).

### Comparison between mCherry- and GFP-CLIP expression vs condensation phenotypes

Cells were plated on 96 well plates and transfected using the TransIT-X2 (Mirus) transfection reagent with 0.075 µg DNA per well according to the manufacturer’s instructions. Images were acquired using a confocal automated microscope (Molecular Device) with a 60X water objective. The first 100 cells observed for each experiment were binned according to their condensation phenotype, as detailed in Figure S2A and B.

### SDS-PAGE and Western blot

Purified proteins or cell lysates were boiled, diluted into sample buffer containing Coomassie dye and x % SDS, and run on gels containing 10% Agarose. After protein separation, gels were stained with QuickBlue Protein Stain Coomassie dye (Lubio Science) for at least 2 hours before destaining in water.

Cell lysates were boiled run on SDS-PAGE gels (10% acrylamide) and subsequently transferred to a nitrocellulose membrane using an iBLOT 2 Gel Transfer Device (ThermoFisher Scientific, IB21001). Nitrocellulose membranes were blocked for 1 h with 5 % dried milk resuspended in TBS-Tween 1 %, then incubated over-night with primary antibodies: anti-beta-tubulin (Sigma, T6074, 1:1000 dilution) anti-EB3 (ATLAS anti-MAPRE3, HPA-009263, 1:500 dilution), or anti-CLIP-170 (Santa Cruz Biotechnology, SC-28325, 1:1000 dilution). The following day, unbound antibodies were washed off with TBS-Tween 1%, and membranes were incubated with secondary antibodies conjugated to horseradish peroxidase (anti-mouse or anti-rabbit; GE Healthcare 17097199 and 16951542, 1:5000 dilution) for 1 hour at room temperature. Following secondary antibody incubation, membranes were washed extensively with TBS-Tween 1% and imaged using an ECL Western blotting detection kit (Advansta) and with Fusion Solo Vilber Lourmat camera (Witec ag).

### Statistical analysis

Statistical analyses were carried out using GraphPad Prism software v9 as described in figure legends. Unless otherwise noted, analyses were carried out between experimental means using one-way ANOVA Fisher’s LSD test, or two-tailed Student’s t-test. P-values less than 0.05 were considered statistically significant.

